# Reduced function of the RNA export factor, *Nxt1*, in *Drosophila* causes muscle degeneration and lowers expression of genes with long introns and circular RNA

**DOI:** 10.1101/543942

**Authors:** Kevin van der Graaf, Katia Jindrich, Robert Mitchell, Helen White-Cooper

**Affiliations:** School of Biosciences, Cardiff University, United Kingdom

## Abstract

The RNA export pathway is essential for export-competent mRNAs to pass from the nucleus into the cytoplasm, and thus is essential for protein production and normal function of cells. *Drosophila* with partial loss of function of *Nxt1*, a core factor in the pathway, show reduced viability and male and female sterility. The male sterility has previously been shown to be caused by defects in testis-specific gene expression, particularly of genes without introns. Here we describe a specific defect in growth and maintenance of the larval muscles, leading to muscle degeneration in *Nxt1* mutants. RNA-seq revealed reduced expression of many mRNAs, particularly from genes with long introns in *Nxt1* mutant muscles. We further determined that circRNAs derived from these same genes are also reduced in the mutants. Despite this, the degeneration was rescued by increased expression in muscles of a single gene, the costamere component *tn (abba)*. This is the first report of a specific role for the RNA export pathway gene *Nxt1* in muscle integrity. Our data on *Nxt1* links the mRNA export pathway to a global role in expression of mRNA and circRNA from common precursor genes, *in vivo*.

**Author summary:** In eukaryotic cells, the DNA encoding instructions for protein synthesis is located in the nucleus. It is transcribed into pre-mRNA, which is processed at both ends and spliced to remove internal spacer regions (introns) to generate mRNA. This mRNA is then transported to the cytoplasm by the mRNA export pathway via nuclear pores, and then used as a template for protein synthesis. We have previously shown that reduction in activity of a specific protein in the mRNA export pathway, Nxt1, has an additional role in testis-specific transcription. Here we describe a further role for this protein specifically in gene expression, particularly of genes with long introns and circular RNAs, and in muscle maintenance. *Drosophila* larvae with reduced Nxt1 activity have normal muscle pattern when they are small, but show muscular atrophy and degeneration as they grow, resulting in significant defects in their movement speed. We discovered that expression of many genes is reduced in the *Nxt1* mutant larvae, but that restoring the expression of just one of these, *abba*, the *Drosophila* homologue of *Trim32* (a human gene involved in muscular dystrophy) is capable of preventing the muscle degeneration. Our data provide a new insight into how defects in generally acting cellular factors can lead to specific defects similar to human diseases.

## Introduction

The formation, growth and maintenance of the musculature is critical for normal animal function, and defects in these processes can lead to impaired mobility, shorter lifespan or early lethality. The somatic muscular tissue of *Drosophila melanogaster* is generated via a highly regulated developmental programme in embryogenesis, such that the first instar larva contains a stereotyped pattern of muscles [1]. The abdominal A2-A7 segments each contain 30 muscles on each side of the animal, each with defined size, shape, position and attachment sites. Each muscle is a single multinucleated cell, derived from fusion of muscle founder cells with fusion competent myoblasts (reviewed in [2]). Very severe defects in embryonic myogenesis result in a failure of the embryo to hatch; less severe defects in the development of the muscle pattern or in muscle function can result in animals with impaired mobility. During larval stages, no new cells are added but the muscles grow extensively by addition of new myofibrils, particularly during the final larval instar (reviewed in [3]).

Normal muscle function is essential during pupariation, and defects in larval muscles can lead to lethality at this stage. About 12 hours before pupariation, a pulse of ecdysone triggers the larvae to stop feeding and move out of the food. About 6-10 hours later they stop moving, contract longitudinally, evert their spiracles and become a white pre-pupa. Three hours later the prepupa contracts from the anterior partially withdrawing the anterior tracheal lining [4]. At this stage, an air bubble forms in the abdominal cavity. The bubble gradually increases in size and the abdominal tissue is forced against the body wall. One hour later, the prepupa becomes separated from the puparium due to the secretion of the prepupal cuticle. The air bubble migrates to the anterior by abdominal muscular contractions, creating space inside the puparium to evert the head, which up to this point has been developing internally [4]. Orchestration of these events requires pristine muscle function.

Regulation of gene expression is important for the normal development and functioning of organelles, cells, tissues and the whole organism. Regulation of mRNA expression depends on transcriptional regulatory processes, such as transcription factor and repressor binding, that influence the ability of RNA polymerase to bind to the promoter site and carry out transcription. Post-transcriptional regulation of RNA adds additional layers of potential control, both in the nucleus and after mRNA export to the cytoplasm. Within the nucleus, critical processing and control points include (alternative) splicing of exons and association of the RNA with mRNA nuclear export factors. The passage of the mRNA transcripts to the cytoplasm requires the export factors associated with the transcript to be recognized by receptors in the interior of the pores (reviewed in [5]). In *D. melanogaster*, the association of nascent RNA with export factors occurs co-transcriptionally and involves first the binding of hnRNPs to the transcript, then recruitment of the THO export complex, and finally recruitment of a heterodimer of Nxf1/Nxt1. The THO complex interacts with UAP56 and REF to form transcription-export complex (TREX) [6]. During the recruitment of Nxf1/Nxt1, export factors such as UAP56 and THO complex are displaced from the RNA. Splicing of pre-mRNAs not only results in removal of the intron sequence from the transcript, but also results in the deposition of a protein complex, the Exon Junction Complex (EJC) 5’ of the splice junction. The EJC has many regulatory roles, including acting to help recruit the RNA export components, and thus facilitates nuclear export of correctly processed mRNAs (reviewed in [7]).

While most pre-mRNAs are spliced to generate linear mRNA molecules, a small proportion of generate circular RNAs (circRNAs) by splicosome-dependent back splicing of segments of primary transcripts (reviewed in [8, 9]). The 3’ end of an exon splices to the 5’ end of the same exon, or to a further upstream splice acceptor site in some cases. The production of circRNA depends on alternative transcript processing; typically, the exon that is circularized is not subject to conventional alternative splicing, but is flanked by relatively long introns [10]. Modulation of transcription rate and splicing rate both affect the relative efficiency of forward and back splicing events, and thus can alter the expression level or ratio of the alternative products [11, 12]. Once formed, circRNAs are relatively stable, and, while typically of relatively low abundance, they can accumulate to levels equal to or even exceeding that of the mRNA(s) derived from the same gene. Analysis of total RNA sequencing data has revealed at least 2500 circRNAs expressed in *Drosophila melanogaster* [13]. CircRNAs are particularly abundant in neural tissue, but all analysed tissues express some circRNAs [13]. CircRNAs have been shown to have a variety of roles *in vivo*, including acting as miRNA sponges, protein sponges, protein scaffolders and being translated (reviewed in [8, 9]).

Human muscular dystrophies are a group of inherited genetic conditions that cause muscles to weaken, leading to an increasing level of disability. Some, such as Duchenne muscular dystrophy and limb girdle muscular dystrophy 2H, are caused by mutations in genes encoding muscle structural proteins (*dystrophin* and *Trim32* respectively) [14, 15], while others, like myotonic dystrophy, are caused by defects in RNA metabolism (Reviewed in [16, 17]). For example DM1 myotonic dystrophy, a trinucleotide repeat in the 3’ UTR of the *DMPK* gene results in sequestration of the splicing regulator MBNL1, and this in turn causes defects in muscle gene expression, particularly alteration of splicing patterns. Interestingly, both MBNL1 and *muscleblind* (*mbl*), the *Drosophila* orthologue of MBNL1, produce abundant circular RNAs [11]. Moreover, Mbl protein is implicated in regulation of production of its own circRNA [11], although the MBNL1 circRNA has not yet been implicated in myotonic dystrophy [18].

We have previously described a role for Nxt1 in regulation of transcription in *Drosophila* testes, indicating that the level of this gene product is critical for processes beyond its known role in mRNA nuclear export [19]. Here we describe a specific role for Nxt1 in the maintenance of larval muscles. Specifically, *Nxt1* partial loss of function animals showed muscle degeneration during the extensive growth associated with the final larval instar stage. We demonstrate that this is probably due to its role in the RNA export pathway as knock down of other RNA export factors caused a similar muscle degeneration phenotype. We found that normal expression of many genes, particularly those with long introns that are sources of circRNAs, requires *Nxt1*. We discovered that both the mRNA and circRNA products of these *Nxt1*-responsive genes are reduced in mutants, although the nascent transcripts of these genes are produced at normal levels. Despite the large number of altered transcripts we were able to rescue the muscle degeneration of *Nxt1* mutants, but not the pupal lethality, by expression of the *Drosophila* homologue of *Trim32*, *tn (abba)*, in muscles. Together, our data indicate an unexpected role for *Nxt1* in post-transcriptional regulation of many genes and circRNA splice products, and show that this RNA export protein is essential for muscle maintenance.

## Results

### Nxt1 pupae have a distinctive curved shape, uneverted spiracles and fail head eversion

We have previously described that *Nxt1^z2-0488^ / Nxt1^DG05102^* transheterozygotes are male and female sterile, but also have significantly reduced viability [19]. *Nxt1^DG05102^* is a null allele caused by a P-element insertion into the coding sequence of the gene, while *Nxt1^z2-0488^* is a hypomorphic allele that disrupts a hydrogen bonding network in the core of the protein thus reducing protein stability [19]. Many *Nxt1^z2-0488^ / Nxt1^DG05102^* transheterozygote pupae had a curved shape and uneverted spiracles (Figure 1A). To quantify this morphological defect we measured the axial ratios from the pupa by measuring the length (excluding the posterior and anterior spiracles) and width. This confirmed that the mutant pupae were significantly longer and thinner than wild type (t-test, p= 1e-08). Scanning electron microscopy (SEM) revealed that the surface structure of the pupal case was similar between mutant and wild type (Figure 1C-E) while in Nxt1 trans-heterozygotes, 50% larvae (N=36) had uneverted anterior spiracles (Figure 1E’’).

**Figure 1.**
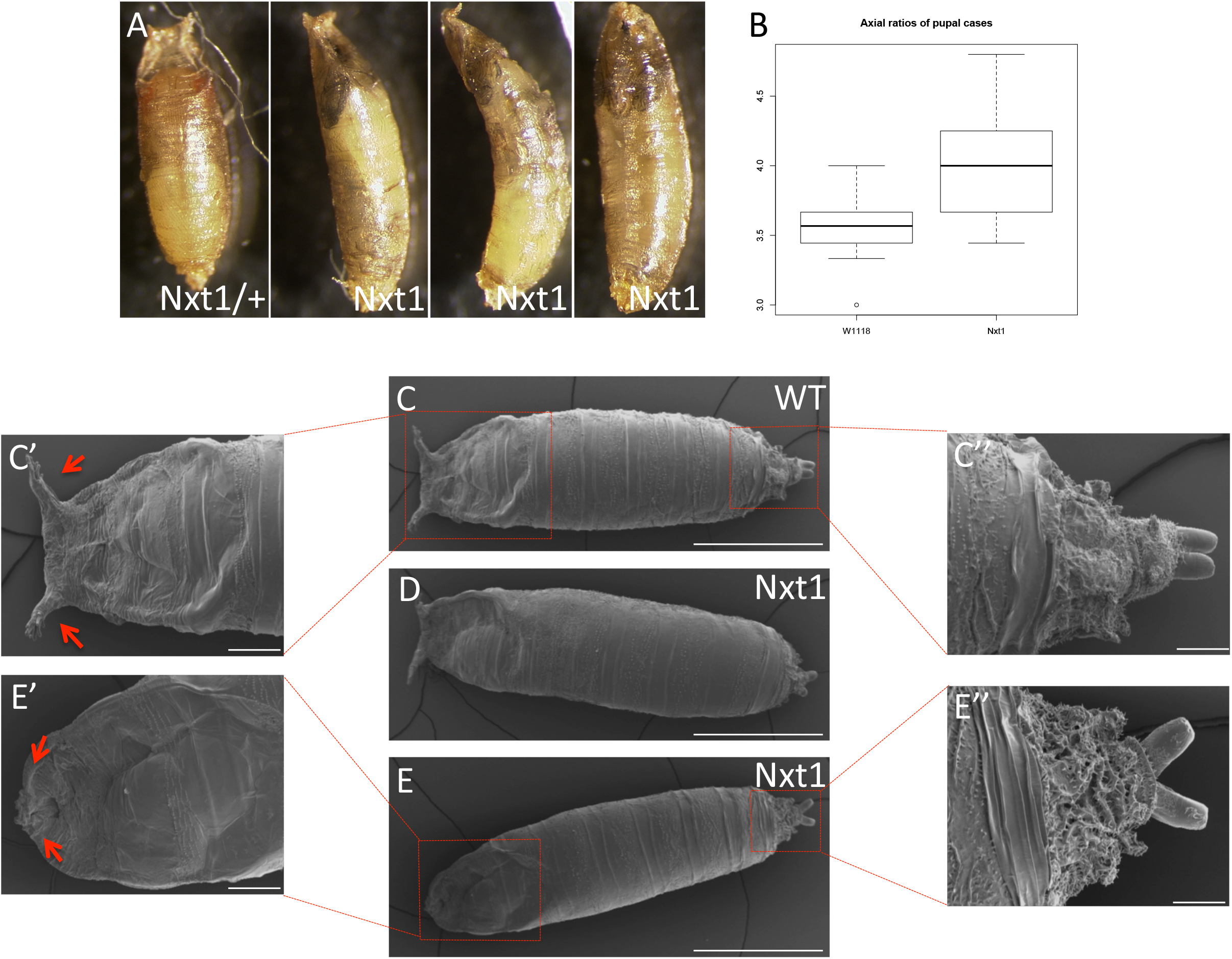
Morphological defects in Nxt1 trans-heterozygote pupae. A) Nxt1 mutant pupae are typically thinner than wild type, are curved, and have uneverted spiracles. B) Axial ratios show Nxt1 mutants are thinner and longer than wild type. C-E) Scanning electron microscopy shows details of anterior and posterior spiracles wild type and Nxt1 mutant pupal cases. Figure in A was taken from [19].

We have also previously reported that the *Nxt1^z2-0488^ / Nxt1^DG05102^* transheterozygote pupae show failure in head eversion [19]. Air bubble formation and migration is implicated in head eversion and therefore the viability of the pupa. To analyse this process we filmed *w^1118^* and *Nxt1^z2-0488^ / Nxt1 ^DG05102^* trans-heterozygote pre-pupae and for 15 hours to observe the air bubble; a typical series is shown. (Supplementary Figure 1) In wild type controls, head eversion occurs at around 12 hours into the pupa phase (Supplementary Figure 1M). Before the head eversion occurred, there was a dramatic period of muscular contractions, as the posterior of the larva wriggled then contracted towards the anterior. At the same time as head eversion, the whole larva body wriggled back within the pupal case to reach the posterior end. The air bubble formed normally in the mutants, and was faintly visible from 0h after pupa formation (APF), growing in size over the next few hours (Supplementary Figure 1A). However, air bubble migration stalled by 8h APF (Supplementary Figure 1I); it remained in the middle of the pupa until 11h APF (Supplementary Figure 1J-M) before disappearing and leaving an empty space at the posterior of the pupa body (Supplementary Figure 1N-P). No head eversion was seen in this mutant pupa (Supplementary Figure 1M).

Crosses of *Nxt1^DG05102^/CyO, Act-GFP* to *Nxt1^Z2-0488^*/ *CyO, Act-GFP* did not reveal the expected Mendelian ratios of 1:2 (Nxt1 trans-heterozygotes: *CyO, Act-GFP;* note *CyO, Act-GFP* homozygotes are lethal). Instead, the observed ratio was 1:10. We assessed the progression of pupal development at four time points (24h, 48h, 72h and 96h pupa), and the number of viable adults. Only about 20% (N=182) of the Nxt1 trans-heterozygote pupae survived through to adulthood, compared to 90% (N= 287) viability for control *w^1118^* pupae (Supplementary Figure 2). The *Nxt1^DG05102^*/+ showed a slight reduction to pupa viability (60% (N= 211); Supplementary Figure 2). Head eversion occurs roughly 12 hours into the pupa stage. Pupa metamorphosis continued (such as body, bristles and wings) in *Nxt1* trans-heterozygotes where most pupa did not have head eversion. The consequence of no head eversion was seen between 48-96 hours into metamorphosis as development stopped abruptly, and the pupae blackened.

### Ecdysone-responsive gene expression is not affected in *Nxt1* mutants

The air bubble phenotype suggests that there are defects in the function of the larval/pupal abdominal muscles, or in the developmental regulation of their function at this stage. Ecdysone coordinates tissue-specific morphogenetic changes by several pulses throughout the *Drosophila* development (reviewed in [20]). A high concentration pulse is observed immediately preceding the larva-pupa transition, and this pulse is essential to trigger the muscular contractions on pupariation and in the pre-pupa. To understand if a transcriptional defect downstream of ecdysone signalling is responsible for the phenotype, in a manner analogous to the role for Nxt1 in regulation of testis-specific transcripts dependent on the transcriptional activation complex tMAC [19], we performed RNA sequencing of pooled whole larvae before (wandering larvae), during (stationary larvae), and after (white prepupae) the pulse of ecdysone.

We extracted the expression data for known ecdysone-responsive genes [21]. Out of 87 ecdysone-responsive genes, only four were mildly mis-regulated (Supplementary Table 1). Mutants of these four genes do not show an air bubble phenotype [22, 23], therefore, it is unlikely that failure of expression of ecdysone-responsive genes is responsible for the air bubble phenotype in Nxt1 trans-heterozygotes.

### Muscle degeneration occurs in *Nxt1* mutant larvae

To determine whether the defects seen in pupal and pre-pupal stages in *Nxt1* mutants are due to muscular defects, we characterised the structure and function of muscles in the larval stages. A larval movement assay was used to track 1^st^, 2^nd^, and 3^rd^ instar larvae to compare their speed between stages and between *w^1118^* and Nxt1 trans-heterozygotes. Larvae were tracked on an agar plate with a control odour on each side (Figure 2A and B). There was no significant difference in the average speed of mutant larvae compared to wild type at either 1^st^ or 2^nd^ instar stage. Normally, wild type 3^rd^ instar larvae travel up to 5x faster than 2^nd^ instars (Figure 2F). In contrast, Nxt1 trans-heterozygous 3^rd^ instar larvae were significantly slower than control animals (Figure 2C), indeed, they were no faster than 2^nd^ instar larvae (Figure 2G), despite being much larger.

**Figure 2.**
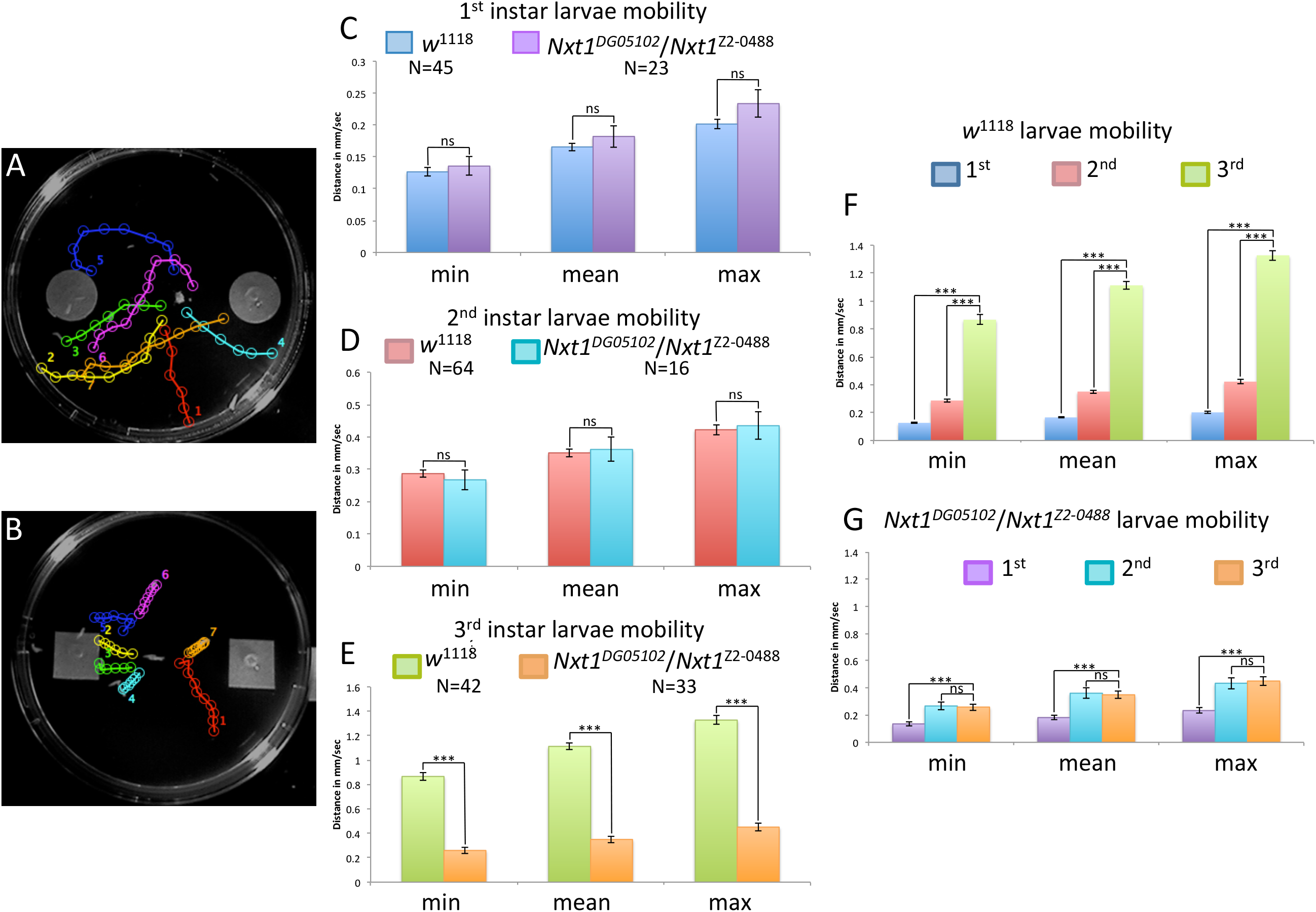
Nxt1 trans-heterozygote 3^rd^ instar larvae have reduced mobility. A-B) Image of agar plate and larval tracks (analysed via MtrackJ) with control substance on the left and odour on the right. C-E) Mobility analysis of 1^st^, 2^nd^ and 3^rd^ instars of wild type and Nxt1 trans-heterozygotes. F-G) All 1^st^, 2^nd^ and 3^rd^ instar data combined for each genotype. Student’s t-test *** p value = <0.001. n.s = not significant.

To investigate the muscle structure in the mutant animals we stained of 3^rd^ instar larval body wall muscles with phalloidin which labels F-actin. In *Drosophila* larvae, each of the abdominal hemisegments A2-A7 has a stereotypical pattern of 30 different muscles [1]. This pattern was clearly observed in the wild type (Figure 3A and C), however, *Nxt1^z2-0488^ / Nxt1^DG05102^* trans-heterozygotes showed clear signs of muscle degeneration (Figure 3B, D and E). Defects seen were variable, and included thinner muscles, fibre splits and torn muscles (Figure 3F-K). We counted the number and nature of defective muscles in 8 hemisegments of 6 control and 6 mutant third instar larvae and categorized the nature of the defects. Defects seen in the mutants were variable and included thinner muscles (15%), loss of sarcomeric structure (22%), degenerating muscles (77%), torn muscles (8%); all control animals had no defects.

**Figure 3.**
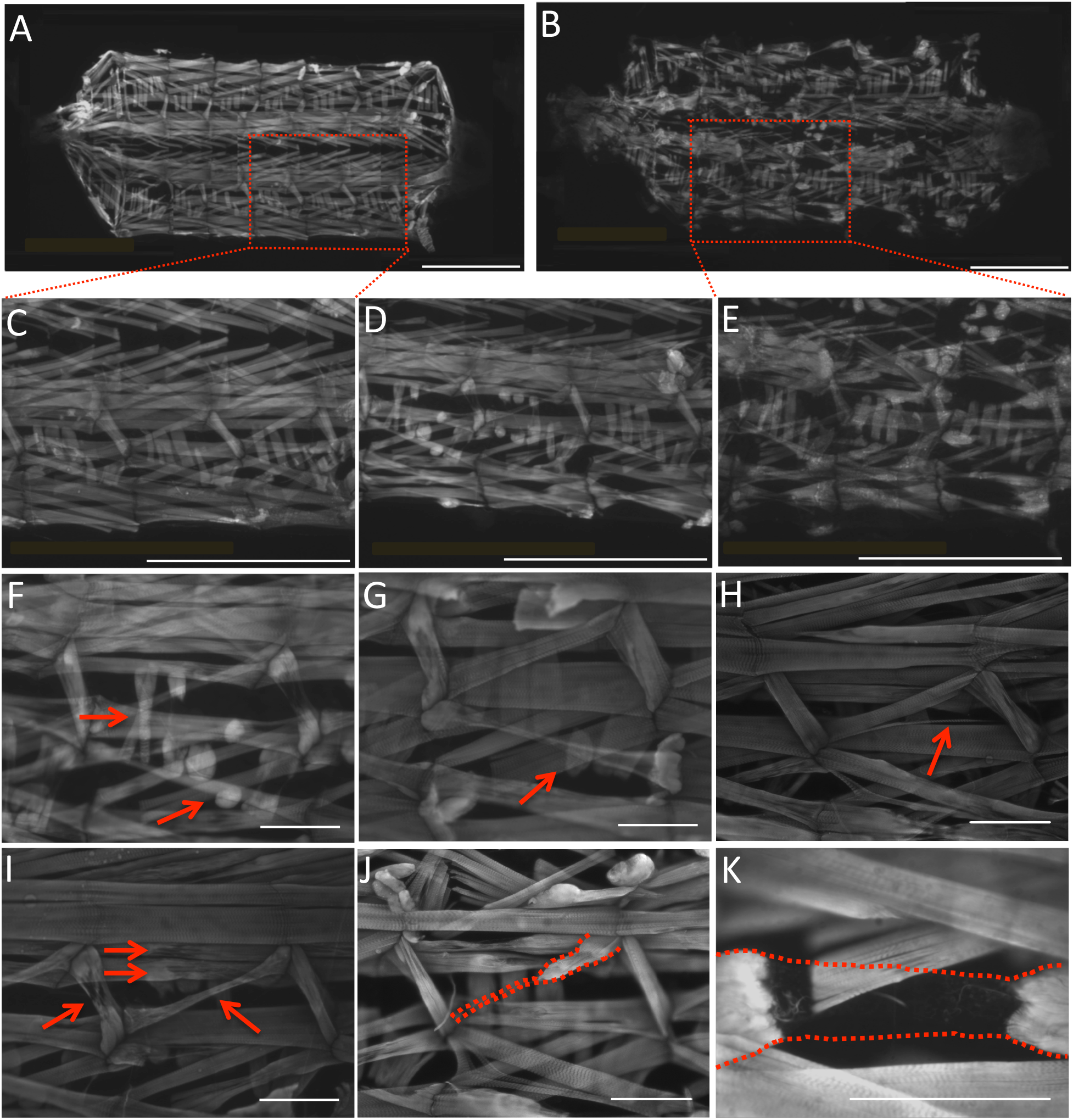
Nxt1 trans-heterozygote 3^rd^ instar larvae show muscle degeneration. Muscles of larval carcass preparations (WT A and C, Nxt1 mutants B and D-K) imaged with FITC-phalloidin. A) Overview of wild type 3^rd^ instar larva muscles. B) Overview of *Nxt1^DG05102^*/*Nxt1^Z2-0488^* 3^rd^ instar larva muscles. C) Higher power image of larva hemisegments from (A). D) Hemisegments from *Nxt1^DG05102^*/*Nxt1^Z2-0488^* with mild muscle degeneration. E) Higher power image of larva hemisegments from (B). F) Lateral Transverse (LT) 1 and 2 crossover (short arrow). F-G) Short LT4 muscle (long arrow). H) Lateral Oblique (LO) 1 fibre split (red arrow). I) J) LO 1 partially degenerated into small bundle of fibres connected to the Segment Border Muscle (SBM). K) Dorsal Oblique (DO) 2 showing strings of fibres (red dotted lines) attached to each end of the muscle. Samples orientated from posterior (left) to anterior (right). A-E) Scale bar = 1 mm. F-K) Scale bar = 200 µm.

In addition to the gross morphological defects, *Nxt1^z2-0488^ / Nxt1^DG05102^* trans-heterozygote mutant animals also sometimes had defects in internal muscle structure. Normal muscle sarcomere structure consists of thick and thin filaments, and phalloidin staining of actin reveals this structure by labelling the thin filaments in the sarcomere (Figure 4A). For Nxt1 trans-heterozygotes, the sarcomere structure was compromised in 22% of the muscles examined. These muscles showed more uniform phalloidin staining indicating weak or absent differentiation of thin vs thick filaments (Figure 4C). This muscle degeneration and sarcomere structure defect phenotype was not fully penetrant, and some Nxt1 trans-heterozygotes had more normal sarcomere structure and less obvious muscle degeneration. This is consistent with the finding that about 20% of the mutant third instar larvae are able to develop to adulthood (Figure 4B). We analysed muscles from first (N=8) and second (N=10) instars with phalloidin staining. This revealed that earlier stage larvae have a normal musculature, and thus that the defects are due to degeneration rather than a developmental defect.

**Figure 4.**
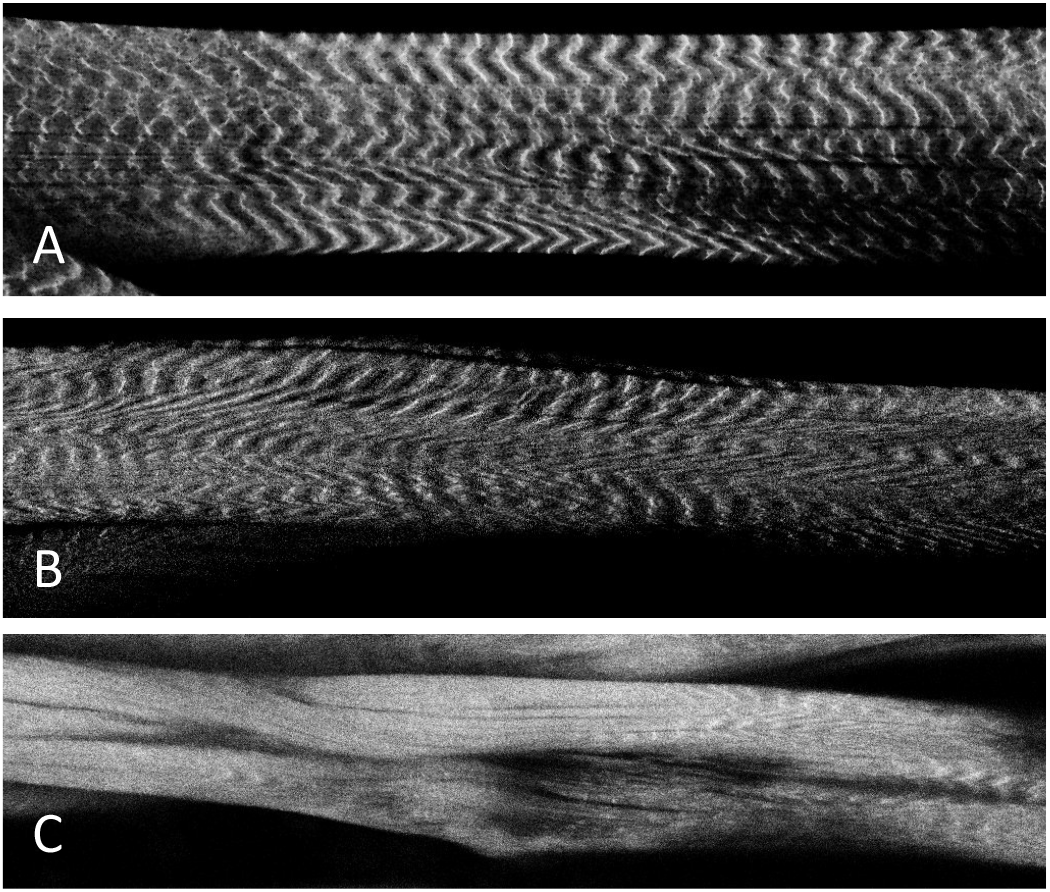
Sarcomere structure compromised in degenerating muscles. A) Wild type muscles stained for actin with phalloidin show normal, ribbed lines, thin filaments. B) Nxt1 trans-heterozygotes non-degenerating muscles with normal, ribbed lines, thin filaments. C) Nxt1 trans-heterozygotes degenerating muscles with compromised thin filaments.

To confirm that the muscle phenotype observed was due to defects in *Nxt1* rather than being a non-specific effect, we used Mef2-Gal4 to drive UAS-RNAi expression specifically in muscles. We combined this with UAS-dicer and high induction temperature to achieve a strong knockdown of the transcript. Both RNAi lines 103146 (chromosome 2) and 52631 (chromosome 3) had previously been shown to effectively knock down *Nxt1* and phenocopy the mutant phenotype in spermatocytes [19]. Phalloidin staining of these 2^nd^ instar larvae showed extensive muscle degeneration (Figure 5). When the temperature was 25°C for embryonic development, with a shift to 29°C after hatching, line 52631 still induced early 2^nd^ instar lethality, while 103146 gave a weaker phenotype of third instar larval lethality, very similar to the *Nxt1^z2-0488^ / Nxt1^DG05102^* trans-heterozygote hypomorphic condition. This indicates that knock down of *Nxt1* specifically in muscles is able to phenocopy the *Nxt1* mutant situation, and shows that the degeneration is due to reduction in *Nxt1* activity in muscles.

**Figure 5.**
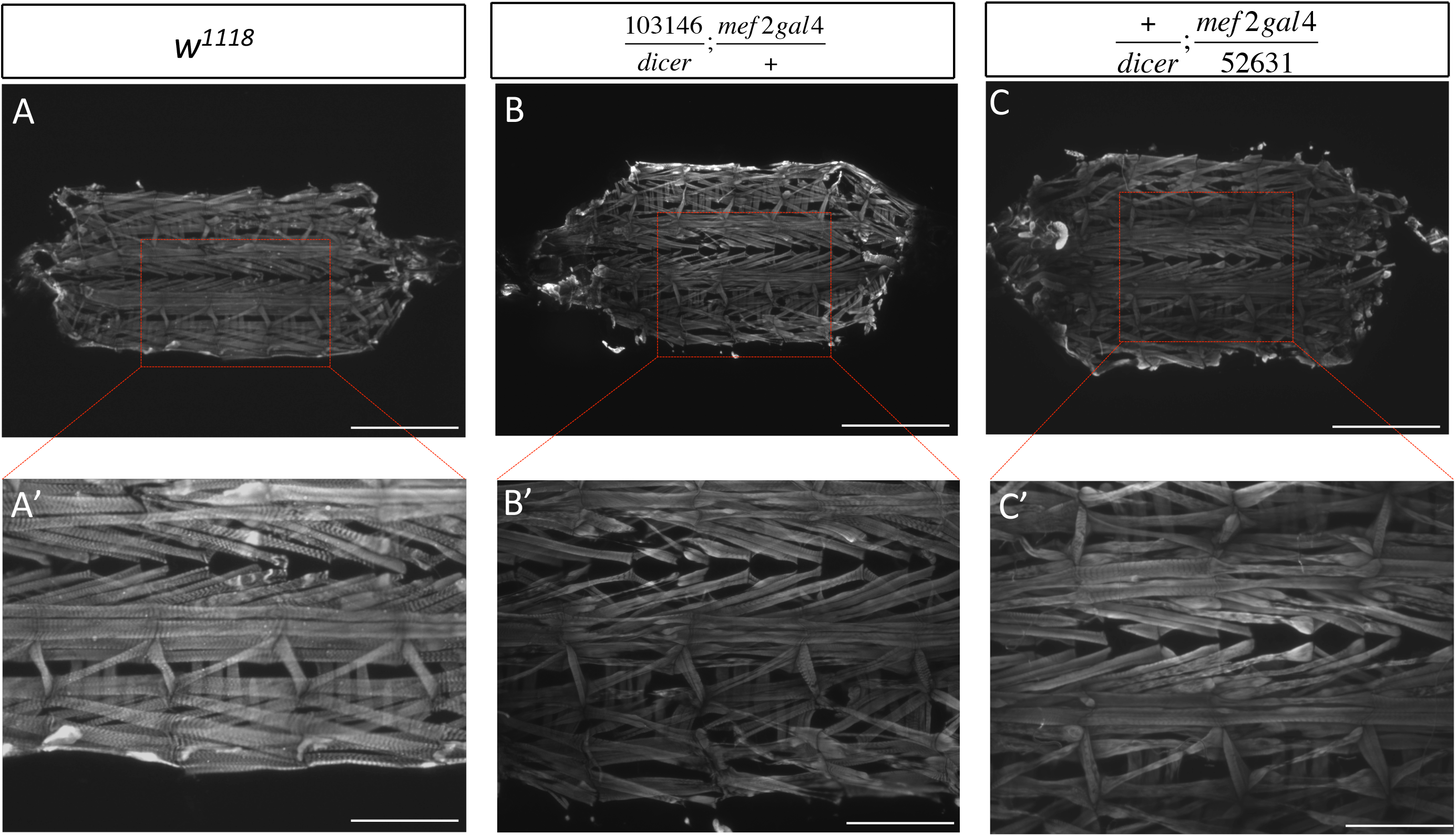
Early 2^nd^ instar muscle degeneration with Nxt1 RNAi. A-A’) Wild type 2^nd^ instar larvae carcass preparations, stained with phalloidin, showing normal muscle composition and no damage. B-B’) Signs of muscle degeneration with the RNAi 103146 line driven by Mef2-Gal4 visible in early 2^nd^ instars with thinner muscles and fiber damage (see insert B’). C-C’) Similar degeneration defects shown for the RNAi 52631 line. A-C) Scale bar = 1mm. A’-C’) Scale bar = 0.5mm.

We expressed GFP-Nxt1 via the UAS-gal4 system both exclusively in muscles at a high level (with mef2-gal4>UAS-GFP-*Nxt1*) and ubiquitously at a lower level (with arm-gal4>UAS-GFP-*Nxt1*) in *Nxt1* trans-heterozygotes. High level, muscle-specific, GFP-Nxt1 expression was able to partially rescue muscle integrity in *Nxt1* trans-heterozygotes (∼13-26% with two 47% outliers (N= 10) Figure 6C) and partially rescue the pupa lethality (∼40% (N= 186); Figure 6A). On average, the axial ratios were similar to wild type, albeit more variable (Figure 6B). Finally, mobility was increased compared to *Nxt1* trans-heterozygotes, but was still significantly reduced compared to wild type (Student’s t-test * p<0.05; Figure 6D). Lower level ubiquitous expression of GFP-*Nxt1* in Nxt1 trans-heterozygotes resulted in improved, but not fully rescued muscle integrity (∼3-33% damage with one 73% outlier (N= 9), increased the pupa viability (∼75% (N= 163)) and increased larval mobility. The axial ratios were similar to wild type (Figure 6B).

**Figure 6.**
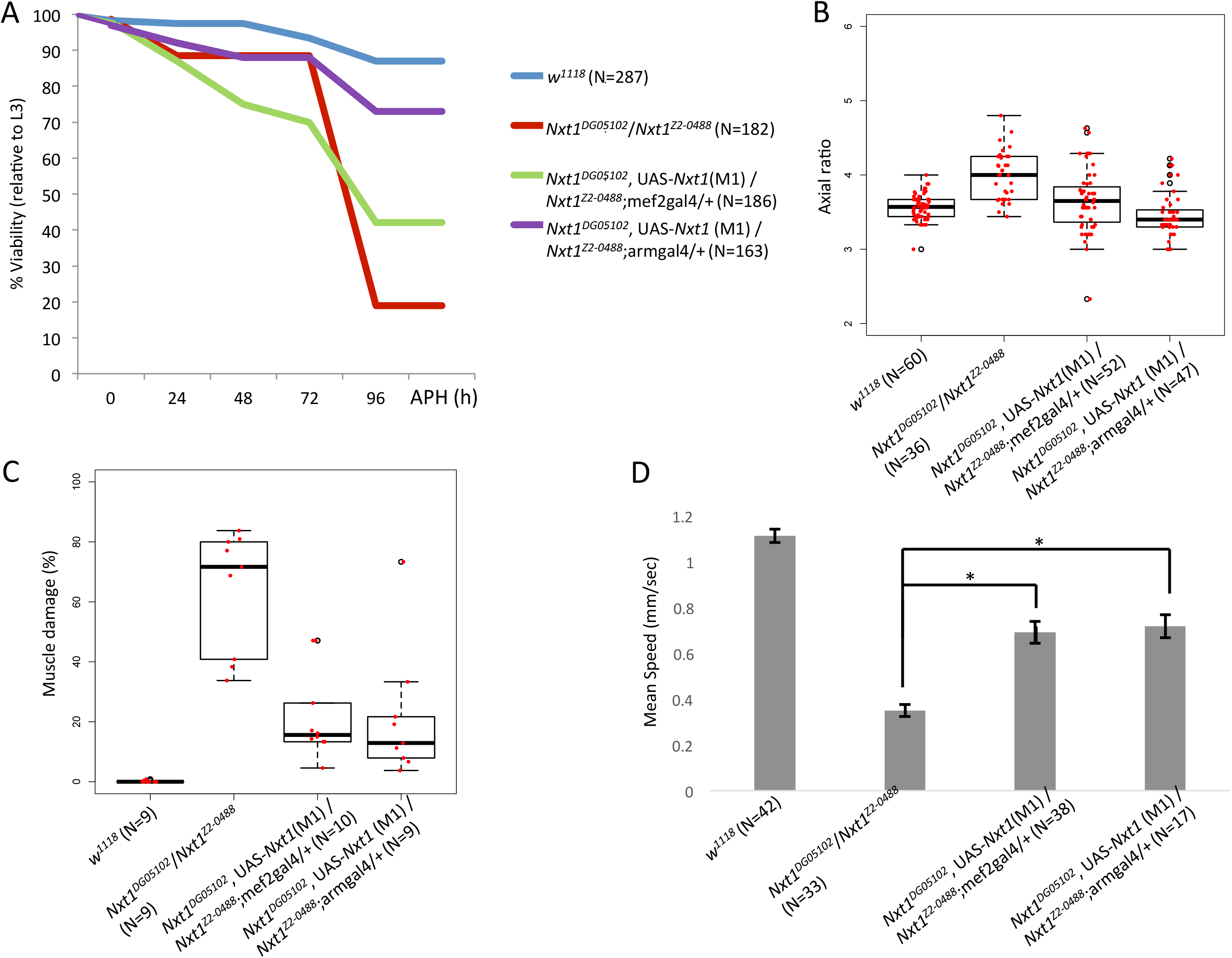
Rescue of Nxt1 trans-heterozygote phenotypes. Nxt1 transheterozygote defects in viability (A), pupal morphology (axial ratios of pupae, B), muscle morphology (C) and 3^rd^ instar motility (D) were partially rescued by over-expression of GFP-Nxt1 in muscles (Mef2-Gal4) or ubiquitously (arm-Gal4).

### Muscle degeneration is associated with larval growth

During the last instar phase, larvae will grow substantially compared to earlier instars. Larvae removed from the food 70 hours after egg laying do not grow, but still crawl until the normal time for pupation (∼112 hours AEL) and may survive through to adulthood, generating very small flies [24]. Larvae at 70 hours AEL are late 2^nd^, or early 3^rd^ instars.

To test whether growth or use (movement) is implicated in the muscle degeneration in *Nxt1* mutants, we fed ∼60 larvae for 70-73 hours then removed them from the food. 14-25% of the larvae (removed from food at 70-73 hr) were able to pupate although less than 5% of those pupae were able to emerge as adults. We simultaneously examined ∼60 of the Nxt1 trans-heterozygote larvae that were starved from 70-hour AEL and found, that only a few larvae pupated and none emerged as adults (similar to *w^1118^*). The pupae that formed had everted spiracles (Figure 7A) and resembled the wild type controls. Interestingly, the larvae that failed to pupate were able to survive for several additional days as larvae before dying. Nxt1 trans-heterozygote larvae were dissected four days after they were removed from the food, and were stained with phalloidin. For all larvae (n=10), no muscle degeneration was observed (Figure 7B). These larvae had been moving normally and thus using their body wall muscles for the four days of starvation. The lack of abnormalities in these animals indicates that the muscle growth, or high levels of force generation associated with third instar larval movement, rather than use per se, is critical for degeneration in *Nxt1* mutants.

**Figure 7.**
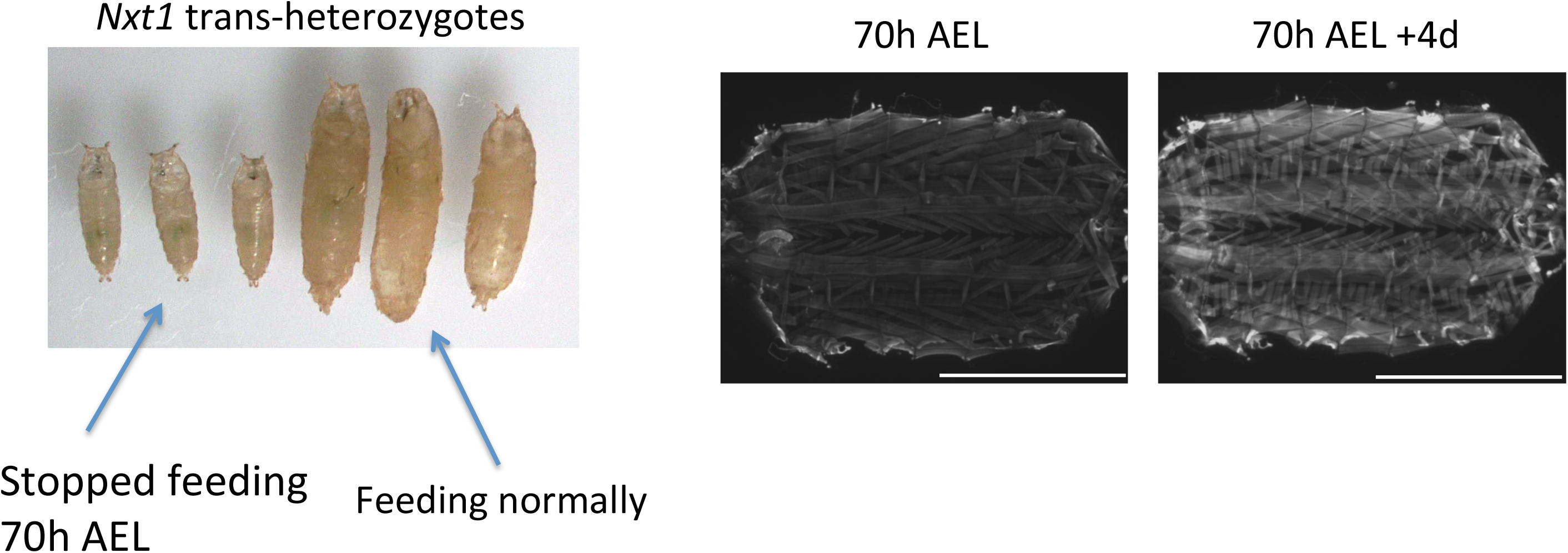
Nxt1 trans-heterozygote muscles do not degenerate when larval growth is prevented. A) Nxt1 trans-heterozygote pupae were smaller, but had normal morphology when food was withheld from 70h AEL. Three food-deprived and three normal feeding animals are shown. B) Nxt1 trans-heterozygote 70h AEL were examined for muscle degeneration. Individuals that had not pupated four days after no feeding still had intact muscles (N= 10).

### Other RNA export factors are also required for maintenance of muscle integrity in *Drosophila* larvae

*Nxt1*’s well characterised function is in the RNA export pathway. To determine whether the muscle degeneration phenotype is caused by defects in this pathway, we assessed the phenotype of knock down of other RNA export pathway genes. We used arm-gal4, to drive RNAi hairpin constructs targeted against *thoc5, sbr (Nxf1), Ref1* and *Hel25E (UAP56)*. This is designed to reduce, but not complete eliminate the RNAi target gene expression throughout the larva. As a positive control we used RNAi against *Nxt1*, and as a negative control we used both *w^1118^* and arm-gal4 alone. RNAi against *Hel25E* caused early larval lethality, even when tested at lower temperature, so we were not able to assess the role of this gene in muscle maintenance. For all the other genotypes, we assayed third instar muscle integrity with phalloidin staining. Knock down of these RNA export pathway components caused a similar phenotype to knock down of *Nxt1*, with muscle thinning and tearing apparent (Figure 8B-E’). Control larvae (arm-gal4 alone and *w^1118^*) had normal musculature (Figure 8A and A’ and data not shown). This indicates that the role of *Nxt1* in muscle maintenance is most likely to be attributable to its role in the RNA export pathway rather than an un-related function.

**Figure 8.**
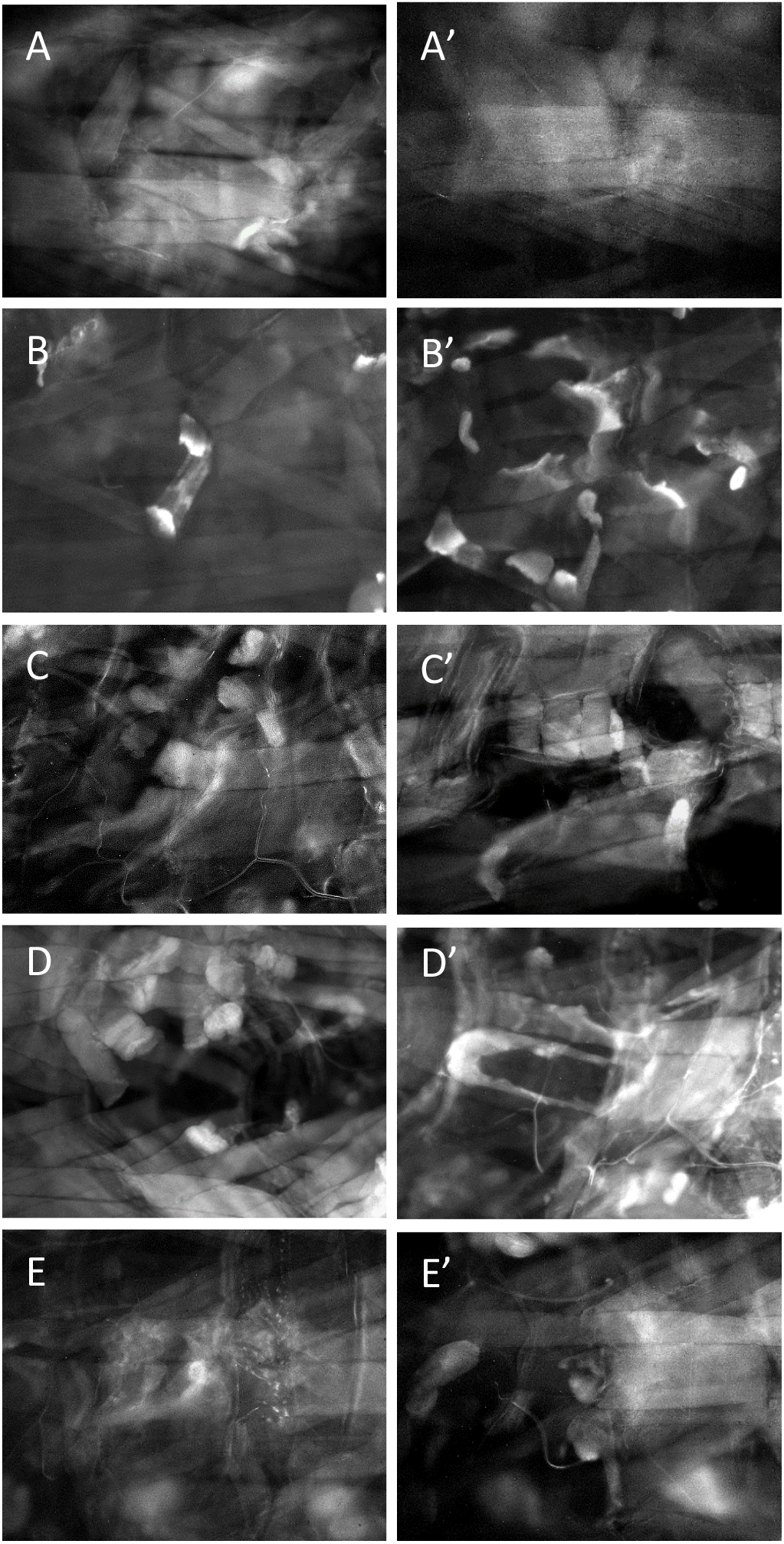
Muscle degeneration caused by RNAi against other RNA export pathway genes. A, A’) Control arm-gal4 driver line late 3^rd^ instar larvae muscles stained with phalloidin shows normal muscle structure. B-E) Muscle degeneration observed after ubiquitous, moderate, induction of RNAi against *Nxt1* (B, B’), *Nxf1* (C, C’), *Ref1* (D, D’) and *thoc5* (E, E’).

### Genes with long introns that also produce circRNAs are sensitive to the loss of Nxt1

The known role of *Nxt1* in RNA export and transcriptional control suggested that the muscle phenotype could be due to transcriptional or post-transcriptional defects in gene expression. To identify somatically expressed differentially expressed genes, we performed RNA sequencing from stationary third instar larval carcasses (comprising primarily muscle and cuticle). Each genotype (*w^1118^* and Nxt1 trans-heterozygotes) was analysed in triplicate. 572 genes were 2x or more up-regulated in mutant compared to wild type, while 1340 genes were similarly down-regulated. The numbers of genes up- and down-regulated at different fold change cut offs are shown in Table 1. To investigate properties of differentially expressed genes we focussed on aspects of RNA processing in the nucleus, and particularly intron length. Nxt1 is important for mRNA export, is recruited indirectly by EJC to spliced transcripts [25], and mutations in the EJC can cause reduced accumulation of transcripts with long introns [26]. In contrast, in *Nxt1* trans-heterozygote testes, transcripts from short and intron-less genes were dramatically reduced. In *Nxt1* trans-heterozygote testes, addition of at least one intron to a down-regulated gene resulted in increased transcript production [19]. We initially analysed the total intron length, number of introns and smallest/largest intron. This revealed that, in direct contrast to the situation in testes, genes down-regulated in *Nxt1* mutant carcass had more introns and a higher total intron length than non-differentially expressed genes, while up-regulated genes had fewer introns and a lower total intron length than the genes that were not differentially expressed (Mann-Whitney test p-value <0.05, except for 16-fold up regulated (p value of 0.16)); (Table 1 and Figure 9). We extended the analysis by also looking at the gene length and shortest/longest transcripts (Table 2). Again, a clear trend showed that down-regulated genes had longer median mRNA length and up-regulated genes had shorter median mRNA length when compared to non-differentially expressed genes (Mann-Whitney p<0.05; Table 2). Finally, the genes down regulated in *Nxt1* mutants produced more mRNA isoforms than those up regulated or not differentially expressed (Table 2). qRT-PCR for 12 genes with long introns confirmed the down-regulation seen in the RNA sequencing data (Figure 10).

**Figure 9.**
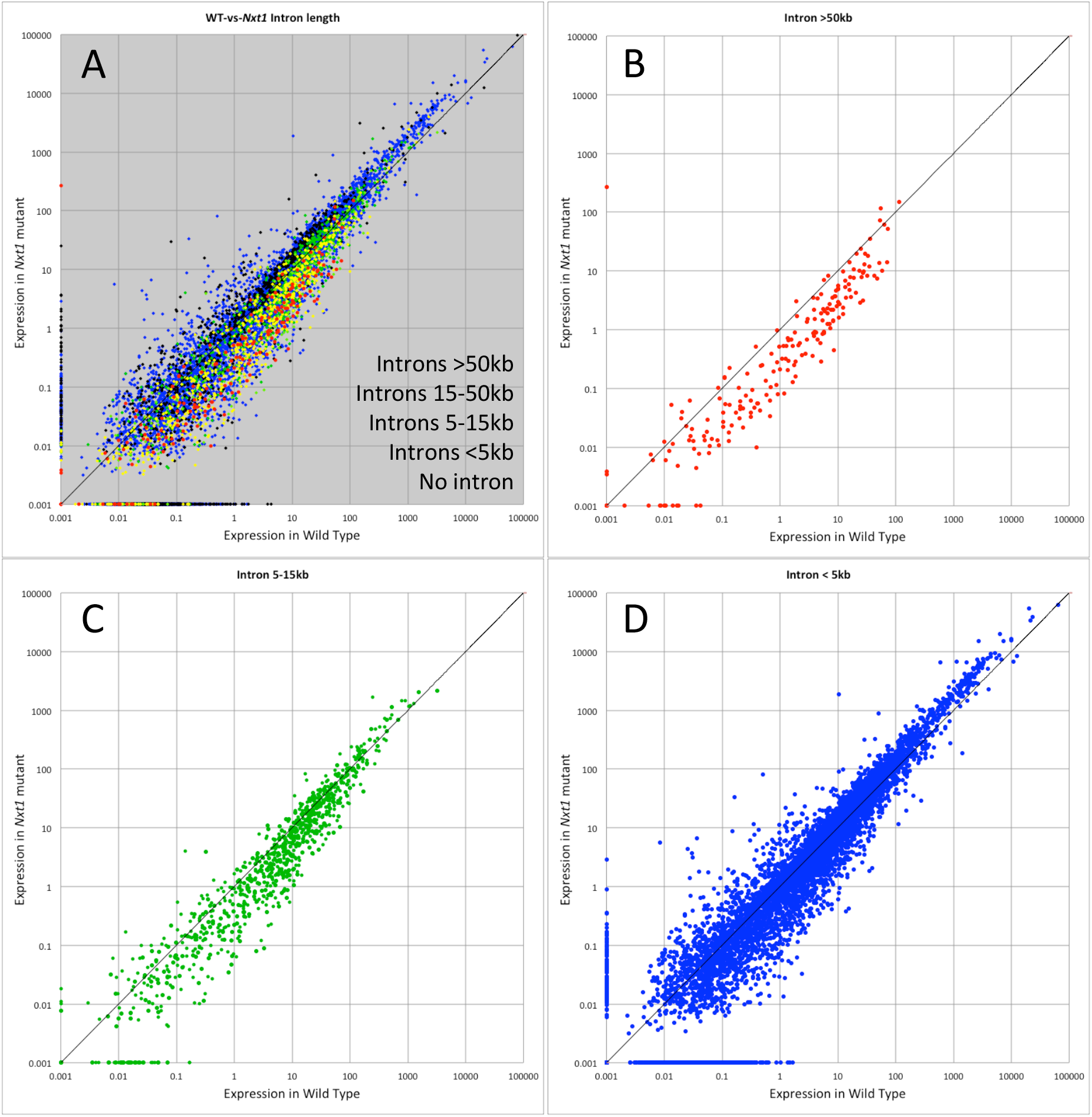
Genes with long introns are under-expressed in Nxt1 trans-heterozygotes. mRNA level (FPKM) showing relative expression in WT and Nxt1 larval carcass samples. A) All genes, coloured to show total intron length. Genes with long introns are typically expressed at lower levels in the mutant than in the wild type sample. B) Genes (red in panel A) whose total intron length is >50kb, C) Genes (green in panel A) whose total intron length is between 5 and 15 kb, D) Genes (blue in panel A) whose total intron length is <5 kb.

**Figure 10.**
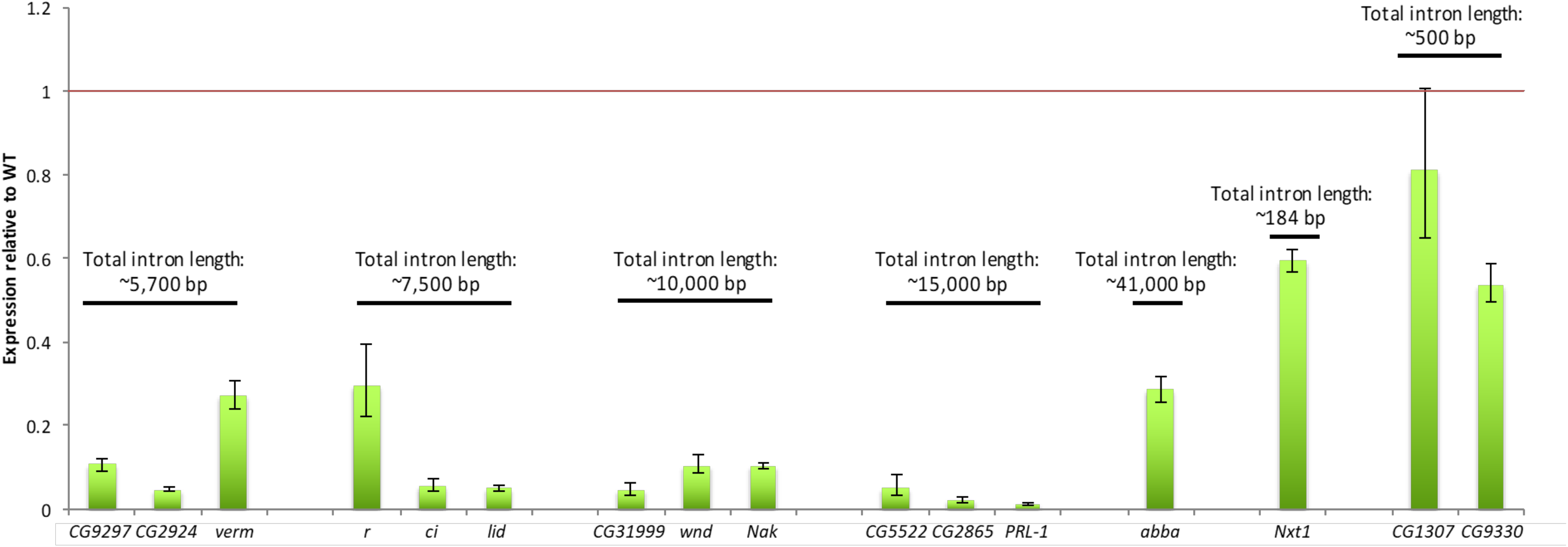
Q-RT-PCR validation of under-expression of genes with long introns in *Nxt1* trans-heterozygote larval carcass. Genes with an average total intron length of 5700, 7500, 10000 and 15000 were examined. Expression of *abba* and *Nxt1* are also shown. CG1307 and CG9330 were used as control genes with low total intron length. All genes examined have expression in the carcass. Rp49 was used for normalization.

**Table 1.** Total intron lengths for genes down and up regulated from larval carcass sequencing. Genes down regulated have longer total intron length compared to up regulated and non-differentially expressed genes. The total intron length becomes longer with the more highly down regulated genes. Down regulated genes have more introns and their largest intron is significantly longer.

| **Median** | **No. of genes** | **Total intron length** | **Number of introns** | **Smallest intron** | **Largest intron** |
| --- | --- | --- | --- | --- | --- |
| Down-regulated >1.5 fold | 1,821 | 5,769 | 9 | 59 | 2,341 |
| Down-regulated >2 fold | 1,340 | 7,594 | 9 | 59 | 2,931 |
| Down-regulated >4 fold | 567 | 10,201 | 11 | 59 | 3,642 |
| Down-regulated >16 fold | 32 | 15,254 | 10 | 63 | 8,081 |
| **Median** | **No. of genes** | **Total intron length** | **Number of introns** | **Smallest intron** | **Largest intron** |
| Up-regulated >1.5 fold | 1,339 | 368 | 3 | 62 | 160 |
| Up-regulated >2 fold | 572 | 352 | 3 | 62 | 155 |
| Up-regulated >4 fold | 89 | 237 | 2 | 62 | 133 |
| Up-regulated >16 fold | 15 | 238 | 2 | 61 | 180 |
| **Median** | **No. of genes** | **Total intron length** | **Number of introns** | **Smallest intron** | **Largest intron** |
| Non-differentiated | 4,754 | 497 | 4 | 60 | 228 |

**Table 2.** Transcript lengths from down and up regulated genes. Genes that are down regulated also have a longer transcript length compared to up regulated and non-differentially expressed genes. For down regulated genes, the number of transcripts is higher and their shortest/longest transcripts are larger.

| **Median** | **No. of genes** | **Gene length** | **No. of transcripts** | **Shortest transcript** | **Longest transcript** |
| --- | --- | --- | --- | --- | --- |
| Down-regulated >1.5 fold | **1,821** | 8,440 | 3 | 3,220 | 4,263 |
| Down-regulated >2 fold | **1,340** | 9,706 | 3 | 3,301 | 4,533 |
| Down-regulated >4 fold | **567** | 12,162 | 3 | 3,722 | 5,168 |
| Down-regulated >16 fold | **32** | 14,213 | 4 | 3,486 | 5,761 |
| **Median** | **No. of genes** | **Gene length** | **No. of transcripts** | **Shortest transcript** | **Longest transcript** |
| Up-regulated >1.5 fold | **1,339** | 1,584 | 2 | 1,034 | 1,186 |
| Up-regulated >2 fold | **572** | 1,411 | 2 | 901 | 1,027 |
| Up-regulated >4 fold | **89** | 1,353 | 1 | 885 | 994 |
| Up-regulated >16 fold | **15** | 1,411 | 1 | 917 | 1,020 |
| **Median** | **No. of genes** | **Gene length** | **No. of transcripts** | **Shortest transcript** | **Longest transcript** |
| Non-differentiated | **4,754** | 2,496 | 2 | 1,642 | 1,856 |

Long introns present a challenge to the spliceosome, and indeed some long introns are spliced as several shorter fragments via recursive splicing [27]. Expression of most genes known to be processed via recursive splicing was detected in our RNA-seq sample (112/116). Only 11 of these were 8x or more down-regulated in the *Nxt1* mutants (34 were 4x or more down-regulated; Figure 9). We concluded that Nxt1 is unlikely to be required for recursive splicing *per se*, and considered other aspects of genes with long introns.

Genes with long introns are sources of circular RNAs, with many circular RNA exons being flanked by long introns [10]. We therefore compared the genes down-regulated in *Nxt1* muscles with lists of genes known to produce circRNA transcripts [13], and found a striking overlap. 466 of the 567 genes 4x or more down-regulated in *Nxt1* muscles had at least one read consistent with a circRNA. 199 of these genes were in the higher confidence set associated with at least 10 circRNA reads. 57 of the 186 genes that were at least 8x down regulated in *Nxt1* muscles were also in the high confidence circRNA set. circRNAs are predominantly produced from genes that are known to have neuronal functions and whose expression is higher in nervous system, even when the analysis is performed on tissues outside the nervous system [13]. Consistent with this, embryonic expression pattern enrichment analysis via FlyMine revealed that the genes down-regulated in *Nxt1* muscles are indeed significantly enriched for expression in nervous-system (such as ventral nerve cord (p=1.13e-8), embryonic brain (p=3.06e-6)). We conclude that Nxt1 is important not only for the export of mRNAs but also for normal expression of mRNAs from genes with long introns, particularly those that produce circRNA products.

### CircRNAs are reduced in *Nxt1* mutants, but nascent transcripts are not

mRNAs and circRNAs are mutually exclusive products made from the same nascent transcript. If Nxt1 and the RNA export pathway controls the ratio between mRNAs and circRNAs, down regulation of Nxt1 would lead to an increase in circRNA products. Alternatively, Nxt1 could be implicated in the stability of the nascent RNA to ensure normal levels of both products, in which case we would expect a reduction in circRNA in target genes in the mutants. In a third model, Nxt1 could act after completion of splicing to stabilise just the mRNA; in this case the circRNA levels would be expected not to change in the mutant compared to wild type. To determine which of these models is most likely, we sequenced both total RNA (after depletion of the ribosomal RNAs) and circRNA (the RNAse R resistant fraction) from larval carcass samples from wild type and *Nxt1* transheterozygotes. The library preparation protocol did not include a selection for poly adenylated transcripts, and thus allowed us to examine pre-mRNA, both spliced and unspliced, as well as circRNA.

We first assessed the levels of circRNAs, focussing only on 336 structures identified in both genotypes with high confidence (backspliced reads identified in at least 2/3 replicates). This revealed that the majority of circRNAs are reduced in expression in *Nxt1* compared to wild type (Figure 11A-D). For these genes, pre-mRNA levels are unchanged in between wild type and mutants, while poly adenylated mRNA levels are reduced in mutants (Figure 11). Similarly, we found no difference between levels of spliced RNAs in when we assessed total RNA samples (Figure 11A-D). Thus, transcription of these genes, and splicing, is not affected in the mutants; however accumulation of poly adenylated transcript is reduced.

**Figure 11.**
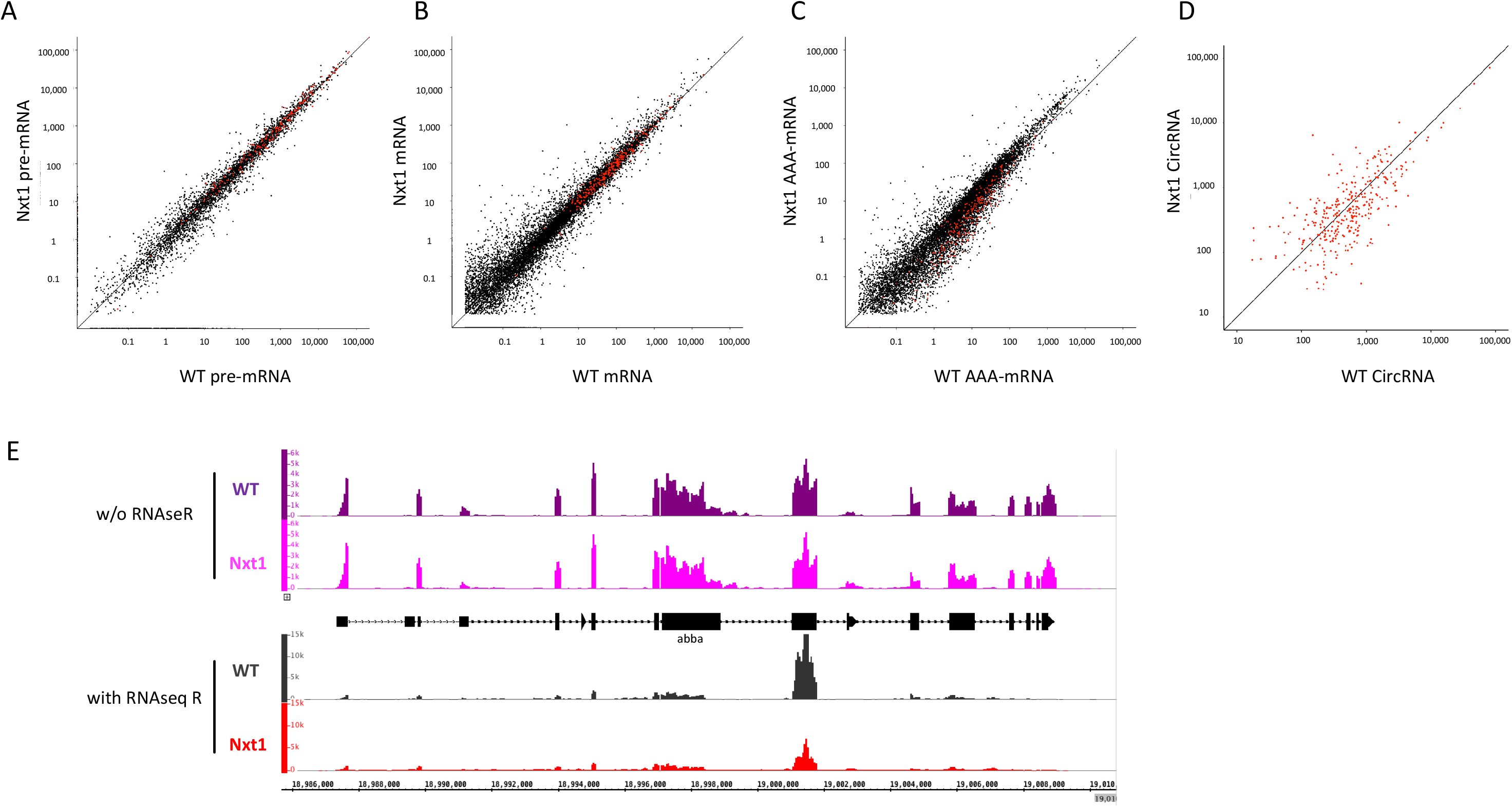
Reduced expression of circRNAs but not nascent transcripts in *Nxt1* trans-heterozygotes. Expression of nascent RNAs (A) and spliced mRNA that is not necessarily polyadenylated (B) is similar in total RNA samples from WT and mutant larval carcasses. Genes that produce circRNAs are highlighted in red. C, sequencing only polyadenylated transcripts reveals reduction in the expression of the mature mRNA from these genes. D, circRNAs detected by sequencing after RNAseR digestion are reduced in mutant larval carcass compared to WT. E, normalised alignment of sequence reads from the *abba* genomic region from the total RNA sequencing (w/o RNAseR) and circRNA sequencing (with RNAseR) reveals that production of a circRNA from this gene is reduced in mutant larval carcasses.

### Increased expression of *abba* rescues the muscle degeneration, but not semi-lethality

Having identified a globlal defect in mRNA and circRNA for many genes, we were interested in understanding how this leads to the muscle degeneration phenotype. The RNA expression of 13 known muscle-specific or muscle-enriched genes was analysed in our original whole larva mRNA-seq data (Supplementary Table 2). Nine genes were more than 1.5-fold up regulated. *abba* (also known as *thin* (*tn*)), the only gene significantly down regulated in this list, is an essential TRIM/RBCC protein that maintains integrity of sarcomeric cytoarchitecture [28]. *abba* loss of function mutations are lethal, with the larvae and pupae being long and thin and having muscle degeneration [29]. *abba* is a large gene with several long introns, which also produces a circRNA derived from circularisation of exon 7 (Figure 11E). The encoded protein has a RING finger, B-Box & CC-domain and several NHL repeats (Figure 12A). qRT-PCR confirmed the reduction in *abba* expression (n=30 larvae per sample; Figure 12). Levels of *abba* mRNA were highly variable between individual stationary larvae (Figure 12) but significantly down in 9 out of 10 larvae, when compared to a pool of 10 WT larvae (Figure 12B). *abba* has at least six isoforms with 13 exons in its longest transcript and contains introns up to 4kb in length. We compared nascent (primers in introns) versus spliced (primers in adjacent exons) abba transcript expression by Q-RT-PCR. Four different regions were selected that would target as many isoforms as possible (Supplementary Figure 3A). Consistent with our total RNA sequencing results, all regions throughout *abba* had similar expression of nascent transcript in mutant compared to control animals, but the levels of spliced transcript were reduced in mutants compared to control (Supplementary Figure 3B). Thus, loss of *Nxt1* does not impact on *abba* transcription, but Nxt1 is important for normal accumulation of the processed mRNA.

**Figure 12.**
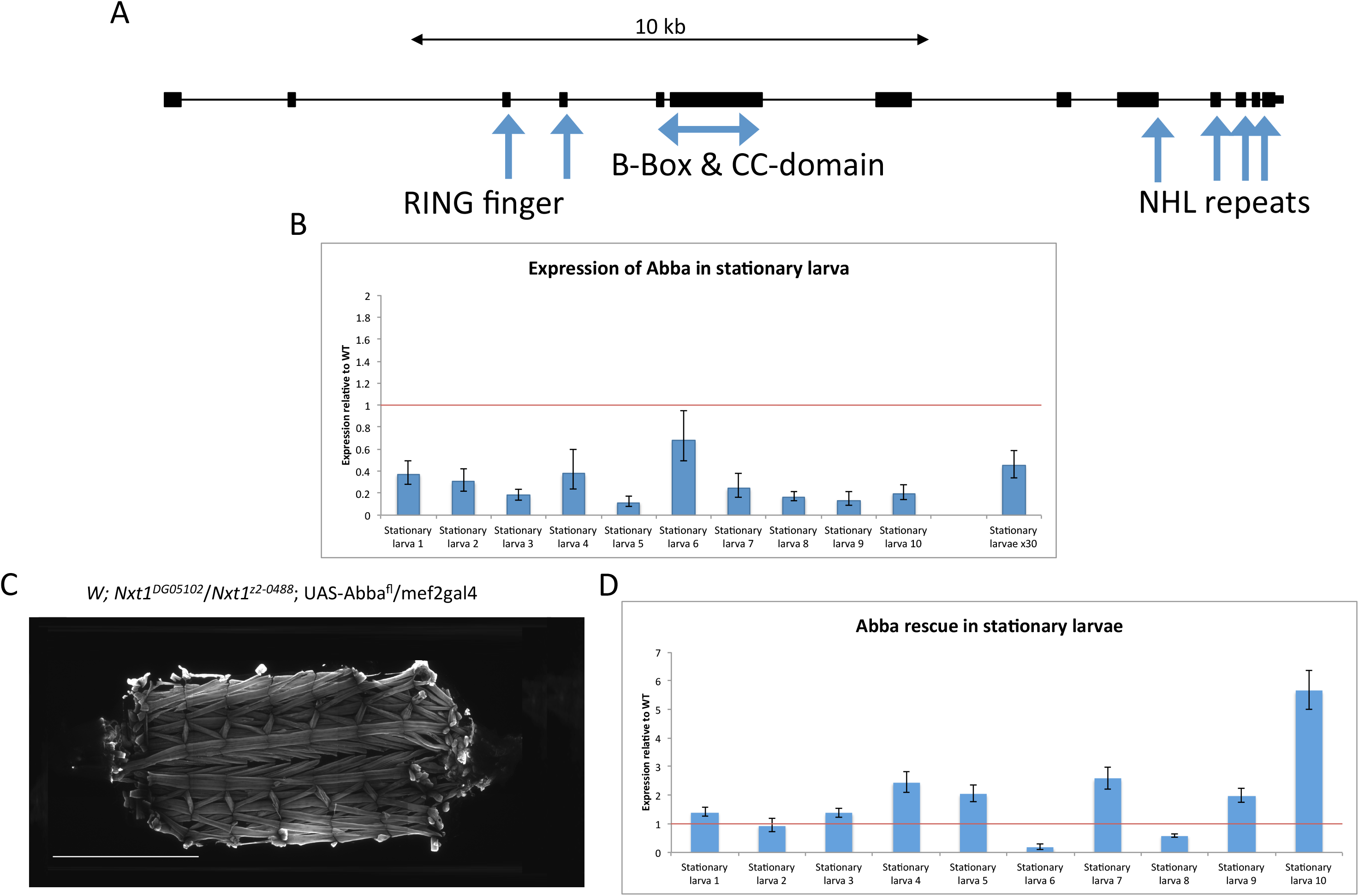
Increasing *abba* expression rescues Nxt1 trans-heterozygote muscle degeneration. A) Genomic structure of the longest *abba* transcript including all known domains, exons are broad boxes, introns are narrow lines. B) Q-RT-PCR analysis of expression of *abba* in 10 individual stationary larvae plus a mix of 30 stationary larvae in Nxt1 trans-heterozygotes, compared to a mix of 10 and 30 stationary wild type larvae, respectively. C) Increasing *abba* with the UAS/Gal4 system specifically in muscles rescues muscle degeneration, as revealed by FITC-phalloidin staining. D) Expression of *abba* in 10 individual larvae from the rescue experiment, compared to a mix of 10 stationary wild type larvae. Rp49 was used for normalization.

To determine whether reduction in the expression of *abba* was implicated in the mutant phenotype, we used a UAS-*abba*^full^ ^length^ ^(fl)^ cDNA construct (kindly gifted by Hanh T. Nguyen; hereafter referred to as UAS-*abba*) and expressed it under the muscle specific driver mef2-gal4 in the *Nxt1* trans-heterozygote background. Staining of the muscles of these larvae with phalloidin, showed that muscle degeneration was fully rescued in 12 out of 16 animals and partially in the other four (Figure 12C). qRT-PCR of individual stationary larvae showed again high variability between individuals, but 8 out of 10 had *abba* expression equal to or exceeding that seen in wild type (Figure 12D).

This result is surprising given the large number of genes whose expression is altered in the mutant larvae. We therefore examined the ability of these animals to survive to adulthood and found that the lethality was not rescued (22%; N=187). These genetic rescue experiments indicate that the muscle defect in *Nxt1* mutant larvae is primarily due to the reduction in *abba* mRNA expression. However, the pupal lethality of most *Nxt1* transheterozygotes is not solely caused by the muscular degeneration, and reduced expression of other target genes is likely associated with reduction in the viability of the animal.

## Discussion

### Reduction in Nxt1 function causes muscle degeneration

Nxt1 is primarily known for its role in the RNA export pathway. Nxt1 binds to Nxf1, which interacts with the nuclear pore complex for exporting mRNA to the cytoplasm [30]. Reducing Nxt1 protein level, reduces the Nxt1-Nxf1 dimer and without Nxt1, Nxf1 interacts less effectively with the nuclear pore complex [31]. Since most mRNAs are exported via the Nxt1-Nxf1 route, it is expected that many transcripts will be affected, and indeed this is what is seen in tissue culture cells [30]. Consistent with this, homozygotes of the null *Nxt1* allele are embryonic lethal. The hypomorphic allele we have used causes reduced protein stability [19], but retains sufficient Nxt1 activity, even when in trans to a null allele, to support mRNA export, and thus allows us to explore other functions of this protein. Intriguingly, the transheterozygotes were able to develop apparently normally to the third instar larval stage, and to undergo pupariation. Most lethality occurred at this transition, with obvious defects in air bubble migration and pupal shape. The phenotype was not fully penetrant, and some Nxt1 trans-heterozygote pupae had normal morphology; 20% of transheterozygote third instar pupae survived through to adulthood.

The defects in pupal morphology were caused by muscle degeneration during the third instar larval stage. Second instar larvae had normal muscles pattern, morphology and mobility. This indicates that the establishment of the larval musculature, which occurs during embryonic development, is not affected by the reduction in *Nxt1* in the hypomorphic allele.

Muscles are composed of tandem arrays of sarcomeres containing thick and thin filaments, arranged in myofibrils. When a muscle twitches, the filaments slide past each other in response to calcium release from the sarcoplasmic reticulum (SR) resulting in force generation. Our examination of the muscle integrity in transheterozygote larvae revealed a variety of defects including muscle atrophy (thinning) and loss of integrity (tearing and splitting). Frequently, we saw loss of filamentous actin in the middle of the muscle while the ends had f-actin; very occasionally we saw balls of muscles associated with loss of attachment or catastrophic failure of muscle integrity and muscle severing. Even when the muscle shape was unaffected, we found a loss of normal internal architecture with disruption of the sarcomeric arrays. Often the muscle was present, but there was filamentous actin staining only towards the ends of the muscle and the central region lacked the sarcomeric structure. It is likely that the reduction in larval mobility, the axial ratio defect, the failure of spiracle eversion and the failure of air bubble movement are all a direct result of the degeneration of muscles in the mutant larvae. All these processes require efficient and coordinated muscle contractions, which are compromised by lack of Nxt1. The curved shape of many mutant pupae is likely to be due to the variability in extent of muscle degeneration: if an individual has more damage on one side than the other, particularly of the longitudinal muscles, there will be unequal contraction as the prepupa forms and one side of the body will shorten more.

During the larval phase, the muscles undergo a ∼50-fold increase in fibre size. This dramatic growth occurs without addition of new cells, as the number of nuclei in the muscle syncytium remains constant [3]. Increased DNA content is achieved via endoreduplication, and increased muscle volume is driven by cell growth associated with new myofibril and sarcomere assembly [3]. The majority of this growth occurs during the third instar larval stage, and depends on nutrient supply and sensing of this via the Insulin/Akt/Tor pathway [32]. Deficits in muscle growth, for example caused by reducing endoreduplication through over expression of cyclinE in muscle, have non-autonomous effects on the growth of the whole larva [32]. We did not detect any difference in the overall size of the third instar mutant larvae, indicating that the defects are not due to defects in nutrient supply or sensing. When *Nxt1* hypomorphic larvae are starved from the late 2^nd^ or early 3^rd^ instar larval stage they remain alive and continue to move, but growth is blocked. This treatment was sufficient to rescue the muscle degeneration that typically occurs during the 3^rd^ instar phase. This indicates that the primary cause of the degeneration is defective (re)-organisation during muscle growth, rather than damage caused by use of the muscles, although we cannot rule out that the higher forces required for third instar larval movement are not also implicated.

### Muscle degeneration can be rescued by increasing *abba*

We were surprised that a hypomorphic allele of a pleiotropic factor such as Nxt1 had such a specific phenotype of muscle degeneration. We reasoned that, while it is likely to be important for the normal expression of many genes, reduction in just one or a few crucial genes could underpin the muscle defects. Our initial RNA-seq of whole larvae was a mixed sex population, and unfortunately the majority of genes differentially expressed between samples were those with testes-specific or testis-enriched expression. This is consistent with our previous findings that *Nxt1* is critical for expression of genes regulated by tMAC in testes [19], but meant that the signal from relatively mild expression changes from somatic tissues was less apparent. Nevertheless, one gene, *abba*, with a known role in muscles stood out as being mildly down-regulated in stationary larvae. The phenotype of *abba* mutants is strikingly similar to that we describe for *Nxt1* hypomorphs, particularly with thinner muscles, long thin pupae and defects in both spiracle eversion and air bubble migration [28, 29]. Abba is a TRIM/RBCC protein involved in maintaining the integrity of sarcomeric cytoarchitecture [28, 29], and is the *Drosophila melanogaster* homologue of human *Trim32*, defects in which cause limb girdle muscular dystrophy 2H [15]. Trim32 is localised to the costamere, which overlies the Z-disk and ensures attachment of the sarcomere to the overlying extracellular matrix via the dystrophin glycoprotein complex.

The phenotypes of defective sarcomere structure, fraying and muscular atrophy are all consistent with reduction in *abba* (costamere) function. The growth of muscles involves the generation of new sarcomeric units, each requiring a new costamere and thus new Abba production and incorporation. The gradual decline in muscle integrity seen during growth may be attributable to reduction in Abba protein levels meaning that the stability of newly formed sarcomeres is reduced. Indeed, in mouse, the expression of Trim32 is upregulated in muscles that are remodelling [33]. Consistent with this model, increasing expression of *abba* using a cDNA construct driven with the Gal4/UAS system is sufficient to rescue the muscle defects seen in *Nxt1* mutants. Starvation from late second or early third instar larval stage is also sufficient to rescue the muscle defect. In this case, there is no muscle growth, so no new sarcomeric units need to be added, the existing structures are maintained, and thus there is no muscle degeneration. The rescue by a cDNA also indicates that it is the reduction in *abba* mRNA, rather than a reduction in *abba* circRNA that is important for the muscle phenotype, as the cDNA construct cannot generate the circRNA.

RNAi constructs to disrupt Nxt1 function in muscles caused degeneration, again without pattern defects even in second instar larvae, indicating that the precise level of Nxt1 function is critical. Very high level, muscle specific, RNAi construct expression, driven from embryonic stages, presumably reduces the level of active Nxt1 in muscles in embryos and early larval stages even beyond that found in the hypomorphic allele combination. In this situation, there would be reduced Abba both during embryonic muscle formation and at the early growth stages, and this would impact on the muscle integrity even before the extensive third instar larval growth period. Driving *Nxt1* RNAi ubiquitously but at a lower level, phenocopies the *Nxt1* transheterozygote situation, with viability to third instar stage and muscle degeneration in these animals. Driving RNAi targeted against other RNA export pathway factors ubiquitously, with arm-gal4, also phenocopies the Nxt1 transheterozygote situation. Importantly, these experiments, coupled with the partial rescue of the muscle defects by expression of Nxt1, confirm that the phenotype is caused by reduction in Nxt1 function rather than by a different gene or by a neomorphic effect of the point mutation in *Nxt1^Z2-0488^*. Additionally, these experiments confirm that the role of Nxt1 in muscle maintenance is due to its function in the RNA export pathway, rather than this being a moonlighting function for the protein.

### *Nxt1* is particularly important for expression of mRNAs and circRNAs from genes with long introns

Transcripts with many and large introns are more sensitive to the loss of Nxt1, which is consistent genes with long introns being sensitive to the loss of the EJC [34]. In direct contrast, genes without introns were particularly sensitive to loss of *Nxt1* in testes [19]. RNA-seq data from larval carcasses, which is highly enriched for larval muscles, allowed us to specifically examine the role of Nxt1 in transcript expression in a somatic tissue in which we have shown its function is critical. This revealed a significant relationship between gene expression in *Nxt1* mutants and intron length for both down and up regulated genes. For down-regulated genes, the more dramatic the down-regulation, the longer the total intron length. Similarly, for up regulated genes, the more up regulated the gene, the shorter the total intron length. No strong changes in EJC component transcripts were found in either whole larvae and larval carcasses. Therefore, it is unlikely that defects in EJC component expression levels are implicated in the phenotype. However clearly the ratio between EJC, Nxf1 and Nxt1 will be affected when the level of active Nxt1 is reduced. During primary transcript processing, the spliceosome removes the intron and then the EJC is recruited to the mRNA [25]. The EJC recruits the THO complex, which also recruits UAP56 and REF to form the TREX complex [35]. UAP56 is displaced by the Nxt1-Nxf1 dimer, binding to REF, then REF is removed from the mRNP and Nxt1-Nxf1 binds directly to the mRNP [36]. This process occurs at every intron, although TREX and Nxt1-Nxf1 can also load onto transcripts in the absence of splicing [37]. Transcripts with many introns have more Nxt1-Nxf1 dimer recruitment. If EJC loading is less efficient on transcripts with long introns, particularly if these are being processed to make circRNAs in addition to mRNAs, then these transcripts could be preferentially sensitive to the Nxt1 trans-heterozygotes where the availability of Nxt1 protein is limiting.

Circular RNAs are a relatively recently described class of RNAs that are produced as alternative products from the same primary transcripts as mRNAs. Global analysis of circRNA abundance reveals that many genes can encode circRNAs, but the propensity for a gene to produce a circular transcript is related to the length of the introns, particularly those that flank the back-spliced exon(s) [10]. Our RNA-seq analysis suggests that the mature mRNAs from genes that also produce circRNAs is reduced in Nxt1 mutants. Additionally, we found that the circRNAs themselves are reduced in the mutants. Mutation of exon junction complex components has also been shown to have a preferential effect on mRNA production from genes with long introns [26]. At least some genes down-regulated in the EJC knock down have been shown to have aberrant splicing patterns in the absence of EJC [26]. In contrast, we found few defects in alternative splicing in *Nxt1* mutants. However, it is also interesting to note that 249/315 genes down-regulated after knock down of the EJC were also on the list of genes that produce at least one circRNA, and 99 are on the higher confidence list of 10 or more circRNAs [13, 26]. The RNA export pathway has already been shown to be linked to 3’ end processing and poly adenylation. TREX subunit THOC5 is recruited to target transcript 3’UTRs by poly adenylation specific factor 100 [38]. Ref association with RNAs is promoted by the 5’ cap, by the EJC, and, crucially, also in the 3’ UTR by nuclear polyA binding protein (PABPN1) [39]. In light of this, it is likely that the EJC and the export adapter Nxt1 (and other factors in the RNA export pathway) are particularly important in processing of transcripts that are alternatively spliced to produce mRNA and circular RNA outputs. We found that pre-mRNA levels of these target genes are not reduced in the mutants, while mature, poly adenlyated mRNAs are, suggesting that the EJC and export factors are acting to ensure stabilisation of the transcripts until completion of 3’ end processing.

RNA regulation, and particularly RNA splicing has been linked to muscular degenerative diseases [17]. Sequestration of the splicing regulator MBNL1 by RNA containing expanded CUG or CCUG repeats reduces functional MBNL and results in aberant splicing of downstream targets in DM patients [16]. The *Drosophila* orthologue of MBNL1, *mbl*, produces an abundant circRNA similarly one of the many MBNL1 splice variants in humans is a circRNA [11]. *mbl* is expressed in both muscles and the nervous system, and is one of the many circRNA-producing genes whose mRNA is reduced (approximately 10x) in *Nxt1* mutant larvae. The production of the *mbl* circRNA is regulated by Mbl protein itself, in an autoregulatory feedback loop. In vertebrates, it appears that MBNL1 does not regulate production of its own circRNA in neuronal cells, but the effect in muscle has not been determined [18, 40]. Our discovery that defects in an RNA export factor, *Nxt1*, can generate a muscular dystrophy phenotype and is associated with differential expression of specific mRNAs and circRNAs in *Drosophila melanogaster* suggests that investigation of the RNA export pathway, and circRNAs, in context of the disease is warranted.

## Materials and Methods

### *Drosophila* culture, strains and genetics

Flies were maintained in standard cornmeal, yeast, dextrose, agar medium, at 25**°**C unless otherwise stated. For selection of staged larvae, 0.05% bromophenol blue was added to the food. Wandering larvae have blue guts, whereas stationary larvae have no blue or very light blue guts. White prepupae were selected by watching larvae, and harvesting them within an hour of pupariation. *w^1118^* was used as a control. *yw; Nxt1^z2-0488^ / CyO Actin-GFP* and *yw; Nxt1 ^DG05102^ / CyO Actin-GFP* were crossed to generate *Nxt1^z2-0488^ / Nxt1 ^DG05102^* transheterozygotes for analyses [19]. The *Nxt1* mutant larvae were selected by the absence of GFP.

VDRC RNAi lines KK107745 (*Nxt1*), P{GD17336}v52631 (*Nxt1*) KK102231 (*thoc5*), KK100882 (*Nxf1*) and KK109076 (*Ref1*), were used to analyse muscle integrity in the third instar larvae [41]. UAS-eGFP-Nxt1 was described in [19].

w;UAS-*abba^fl^*, kindly provided by Hanh T. Nguyen, uses the EST GH06739 (GenBank accession number AY121620) as template to amplify and clone the entire *abba* coding or defined regions of *abba* into the pUAST vector [28]. The 3’ end of the construct encodes an HA tag. UAS-stock is in heterozygous condition. Mef2-Gal4 [42] was kindly provided by Michael Taylor (Cardiff University); Arm-Gal4 and UAS-dicer were from Bloomington stock centre.

### RNA extraction, cDNA synthesis and Quantitative PCR

Total RNA was extracted using Trizol (ThermoFisher Scientific) and then further cleaned up with the RNeasy Mini Kit (Qiagen) according to manufacturer’s protocol. The DNaseI step was included. RNA was quantified with a nanodrop ND-1000 (ThermoFisher Scientific) and stored at −80°C. For larval carcass RNA sequencing, 30 carcasses were used per replicate in triplicates. For qRT-PCR, either a single carcass or a mix of up to 10 carcasses were used as detailed in the results. cDNA was generated using 100ng total RNA and oligo dT primers with the Superscript III kit (Invitrogen). The cDNA reaction was diluted to 60 or 120µl with dH_2_O, and 1µl of this cDNA was further diluted with 7µl dH_2_O to use as a template in the qRT-PCR reactions. 10µl PowerSybr reagent (ABI) with 1µl of a 10µM solution of each target-specific forward and reverse primers (primer sequences on request) were added for a total reaction of 20µl. The qRT-PCR was performed on a Chromo4 instrument (MJR) using the Opticon Monitor 3 software. *Rp49* was used as a control gene for normalization, fold changes were determined by ΔΔCT. All reactions were performed in triplicate.

### Scanning Electron Microscopy

Young pupae (<4h) were picked, cleaned with water, and air dried overnight. Up to 6 pupae were put on an aluminium backing plate of a SC500 sputter target. A BIO-RAD SC500 sputter coater was used to coat the non-conducting pupae with ∼20nm thick layer of Au90Pd10 according to manufacturer’s protocol (Quorum Technologies). After coating, pins were put in a FEI-XL30 Field Emission Gun Environmental Scanning Electron Microscope. For imaging, a 30 μm diameter final aperture with a beam current <1nA was used to take pictures from each pupa at the posterior/anterior end and the middle section.

### Magnetic Chamber Dissection for larval muscle staining

For larval dissections, a magnetic chamber was used [43]. A magnetic strip, with a 30 mm diameter hole in the middle, was glued on a 76 mm x 51 mm slide with 10 mm of the strip sticking out at all sides. Each magnetic chamber uses 2 centre and 4 corner pins. For constructing all pins, see Figure 1 in [43]. All pins were glued to a vintage metal index tab. A larva is put in the middle of the hole with a drop of low Ca^2+^ saline, HL-3 [44]. Larva are put ventral side and two centre pins are used to prevent the larva from moving by pinning at the most anterior and posterior side. Vannas Spring Scissors – 3mm blades (RS-5618; FineScience) were used for cutting. A dorsal incision cut was made at the posterior end with short shallow cuts between the two tracheas until the anterior end was reached.

### Phalloidin staining of larval muscles

The larva was cleaned on the magnetic chamber by removing all internal organs carefully. All Ca^2+^ saline, HL-3 solution was removed with a P-100 pipette and fresh drops were added two more times while cleaning the larval carcass. The larval carcass was fixated by adding fresh 4% paraformaldehyde in PBS for 2 minutes. All the pins were removed from the larval carcass and the sample was transferred to a glass well with 100μl 4% paraformaldehyde for another hour. The fixative was removed, and the larval carcass was washed twice with 100μl PBS-T (0.1% Triton X-100) for 5 minutes each. Two drops of Alexa Fluor^TM^ 488 Phalloidin (ThermoFisher) was added to 1ml of PBS-T. 100μl Alexa Fluor^TM^ 488 Phalloidin mix was added to the well to stain F-actin in larval muscles for 1 hour. Alternatively, FITC phalloidin was used at a final concentration of 1 μg/ml in PBS-T. Phalloidin solution was removed and the larval carcass was washed with PBS-T two times for 5 minutes each. The larva carcass was put on a microscope slide and mounted in 85% glycerol + 2.5% n-propyl gallate. Images were made on an Olympus BX 50 (Olympus) microscope and Hamamatsu ORCA-05G digital camera. For a close-up of the larval muscle sarcomeres pattern, images were taken with a Leica DM6000B upright microscope with HC PL Fluotar 20x/0.50 and HCX PL APO 40x/1.25 oil objectives.

### Criteria for scoring muscle defects in larvae

The integrity of larva muscles was calculated as a percentage from a total of 8 hemisegments. The hemisegments A2-A5 were in the abdominal area and were not damaged by any of the pins. Each hemisegment contained 30 different muscles, so a total of 240 muscles were inspected. The integrity of a muscle is compromised if the muscle is damaged in any way, such as being torn, thin, loss of sarcomeric structure or missing. The muscle damage percentage is the total number of damaged muscles divided by 240 and multiplied by 100.

### Starvation of larvae

Larvae were fed up to 70, 71, 72 and 73 hours after egg laying (AEL) before removal from the food. The larvae were transferred to a petri dish with a moist filter paper in it to prevent desiccation. Larvae in the petri dish were checked at regular intervals to ensure the filter paper was moist and keep track of the progress of the metamorphosis. Larvae deprived of food 70 hours AEL that survived for four days were analysed for muscle integrity.

### Mobility Assay

First, second and third instar larvae were used for mobility analysis. Larvae (up to 10) were put in the middle of a 1% agar dish with 1% paraffin oil on one side and an odour (1% 2-propanol) on the other side. Larvae were filmed with a Samsung SDN-550 camera using the micro manager 1.4 program for 200 frames, with 1 frame per second (fps) in the dark under red light. Movies were analysed with MtrackJ [45] via Fiji. Larvae were tracked per 20 frames (1^st^ instar) or 10 frames (2^nd^ and 3^rd^ instars) for at least 100 seconds (1^st^ instar) or 50 seconds (2^nd^ and 3^rd^ instar) while larvae were continuously crawling on the plate. Mean speed is calculated in mm/sec.

### Viability Assay

Third instar larvae were taken from the standard vial and put in a new vial with a low quantity of fresh food for the viability assay. After 5 days of incubation at 25°C vials were taken out and pupa lethality was backtracked by looking at 6 different points: 1) larvae that did not pupate (0h APF), 2) no visible development in pupae (24h APF), 3) head eversion and development of the eye (48h APF), 4) bristles on dorsal thorax (72h APF), 5) complete fly development in pupa (72h APF) and 6) emerging adults.

### Whole larvae mRNA sequencing and analysis

Eighteen libraries were prepared from total RNA of *w^1118^* and Nxt1 trans-heterozygotes genotypes. Each genotype had samples of three different stages (wandering larvae, stationary larvae and white prepupae) in triplicates. RNA extraction was performed using Trizol (ThermoFisher Scientific) followed by the RNeasy Mini Kit (Qiagen). The samples were sent to the University of Exeter to perform the library preparation (ScriptSeq RNA-Seq Library Preparation Kit (Illumina)) and sequencing. All samples were 100bp paired-end sequenced on an Illumina HiSeq 2500 using standard mode.

Each sample was prepared in triplicate, and a sequence depth between 6.4M – 11.1M was achieved with sequencing all libraries. The data was analysed with the Tuxedo suite [46] via GenePattern browser [47]. The lists of genes and their Fragments per Kilobase of Exon per Million Mapped Fragments (FPKM) values were compared and statistically tested between the genotypes with Cuffdiff and were imported to excel and divided into more than 2-fold down regulated, more than 2-fold up regulated and non-differentially expressed lists. The differentially expressed genes that were assigned statistically significant had a p-value of <0.05. For analysis of ecdysone-responsive genes we looked for differential expression between mutant and control, minimum of 2-fold up or down regulated and a minimum FPKM value of 10 in at least one condition

### Larval carcass mRNA sequencing

Six libraries were prepared from total RNA of *w^1118^* and Nxt1 trans-heterozygotes genotypes. Stationary larvae from each genotype were taken and samples were generated in triplicates. RNA extraction was performed using a combination of Trizol (ThermoFisher Scientific) and the RNeasy Mini Kit (Qiagen). Libraries were generated using the TruSeq Stranded mRNA Library Prep (Illumina) according to manufacturer’s protocol. This library preparation method includes an oligo dT selection to sequence only mature, polyadenylated mRNAs. Samples were sequenced using 2×75bp paired-end with the NextSeq 500/550 Mid Output v2 kit (150 cycles; Cat. No. FC-404-2001) on an Illumina NextSeq500 Sequencer (Illumina). Sequencing was performed by the Genome Research Hub at Cardiff University, School of Biosciences

Each library achieved a read depth between 21M to 32M reads and the data was analysed through the Tuxedo suite [46]. The list of genes with FPKM values provided by Cuffdiff was used to separate the lists in more than 1.5-, 2-, 4-and 16-fold down regulated, more than 1.5-, 2-, 4- and 16-fold up regulated and non-differentially expressed. The differentially expressed genes that were assigned statistically significant had a p-value of <0.05. For graphical display using Excel on log axes, 0.001 was added to all FPKM values.

### Larval carcass total RNA and circRNA sequencing

For total RNA and circRNA sequencing from larval carcass, a total of 30 larval muscle preparations were used for each replicate. Stationary stage third instar larvae were dissected in PBS using the magnetic chamber before being washed twice in PBS, and then transferred directly into an Eppendorf containing 100μl lysis buffer. Batches of 5-7 larval carcasses were pooled in each Eppendorf before being transferred to −20 C until all sample collection was complete. No more than 1hr elapsed between dissection and lysis. *yw; Nxt1^z2-0488^ / Nxt1^DG0510^* larvae were used for mutant analysis, with *w^1118^* controls for comparison.

RNA was extracted using the RNAeasy Mini Kit (Qiagen) according to manufacturer’s recommendation with on-column DNAseI digestion. RNA quality was checked on a High Sensitivity tape on the TapeStation 2200 (Agilent Technologies) and quantity was checked by Qubit. 1 μg of RNA was treated with 20 units of RNAseR (Epicentre) in a 25μl reaction at 37°C for 30 minutes. Mock samples were treated with the same volume of water. All samples were then cleaned up with 2 volumes of RNA Clean XP beads (Agencourt) and eluted with 13 μl of water. 10 μl of mock and RNAseR treated samples were used to make RNA libraries using the TruSeq Stranded total RNA-seq with Ribozero gold LT kit (Illumina) following manufacturers recommendations. Libraries were pooled in equimolar proportions and sequenced on a 2×75bp High-output NextSeq500 cartridge, yielding on average of 40 million reads per sample. All sequencing was performed by the Genome Research Hub at Cardiff University, School of Biosciences.

### Total RNA-seq Analysis

Quality control analyses were performed on all samples using FastQC v 0.10 (http://www.bioinformatics.babraham.ac.uk/projects/fastqc/). Raw reads were trimmed with trimmomatic v.0.36 [Bolger et al, 2014] (HEADCROP:15 LEADING:3 TRAILING:3 SLIDINGWINDOW:4:15 MINLEN:36). Sequence reads were aligned to the *Drosophila melanogaster* genome (Dm6.23) using STAR v.2.5.3a [48] with options: --outMultimapperOrder Random --outSAMmultNmax 1. Flybase r6.23 annotations were supplied to index the genome but annotations were not provided for mapping. A second QC analysis was run using Bamtools v.2.4.1 [49] and the MarkDuplicate tool from the Picard suite v.2.18.14 (https://broadinstitute.github.io/picard/).

High rates of duplicated and multi-mapping reads were observed in all samples. Upon thorough investigation, we concluded that those were not artificial. Duplication rates were due to a very high abundance of a small number of genes, and was linked to multi-mappers (i.e., multi-mapping reads were also duplicated). Multi-mapping was due to a small rRNA contamination, and highly expressed small RNA (e.g. cuticle gene Cpr49Ac) or RNA with repeated conserved domains (e.g. cuticle genes lcp1 and lcp2). Differential expression analyses were also run with and without deduplication and led to the same conclusions. Multi-mappers were excluded from further analyses but duplicated reads were kept.

Read counts were generated using Subread package FeatureCounts v. 1.6.2 [50]; reads mapping to one location were assigned to exons (for mRNA) or introns (for pre-mRNA) and reported by gene_id. (metafeature). DESeq2 (Love et al, 2014) was used to normalise reads counts and calculate fpkm values. For visualization in genome browsers (i.e. Integrative Genomic Browser), strand-specific normalised bigwig files were generated with the bamCoverage tool of the deepyools suite v.3.0.1 (--ignoreDuplicates --effectiveGenomeSize 142573017 --normalizeUsing RPKM).

### CircRNA Identification

Paired-end reads were merged prior to analysis. CircRNAs were identified using PTESFinder v.1 [51] (parameters -s 65 -u) with alignments to the *Drosophila* genome (Dm6.23) and the UCSC Refseq transcriptome. Only those circRNAs that were identified in at least two RNAseR-treated biological replicates were included in further analyses. This is a stringent criterion given the high variability between RNase R-treated replicates and the sequencing depth [52]. 336 circRNA structures were identified in at least two replicates of both WT and *Nxt1* samples and are included in further analyses.

As a measure of circRNA abundance, we used the number of back-splice per million mapped reads (BPM), defined as: # reads supporting each structure generated by PTESFinder / total # mapped reads (junctions+canonical) *10^6. We averaged the two highest biological replicates. The corresponding two mock (non RNAseR treated) replicates were used to quantify the spliced RNA (but not necessarily poly adenylated) and pre-mRNA of the host transcript. Plots were generated in R with the ggplot2 package.

## Acknowledgments

We gratefully acknowledge all members of the Cardiff University *Drosophila* community, and particularly Sonia Lopez de Quinto, Michael Taylor and Wynand van der Goes van Naters for helpful input on experimental design and data interpretation throughout the project. We thank Nick Kent and Angela Marchbank in the Cardiff University School of Biosciences Genomics Hub for help with the sequencing, we also thank Lorna Harries, Karen Moore and Ryan Ames in Exeter University for guidance with the circRNA sequencing protocol and analysis. *Drosophila* stocks were provided by Michael Taylor (Cardiff University), Hanh T. Nguyen (Friedrich-Alexander University of Erlangen-Nürnberg, Erlangen, Germany), Vienna Drosophila Resource Centre and Bloomington Drosophila Stock Centre. RNA-seq was carried out by Exeter University sequencing service and by the Cardiff University School of Biosciences Genomics Hub.

## Data availability

RNA-seq datasets for *w1118* and *Nxt1* transheterozygotes larvae are available via GEO: whole larvae before, during and after puparium formation (GSE125781), and late third instar larval carcass samples (mRNA: GSE125776; total RNA and CircRNA: GSE135591).

---

**Supplementary figure 1.**
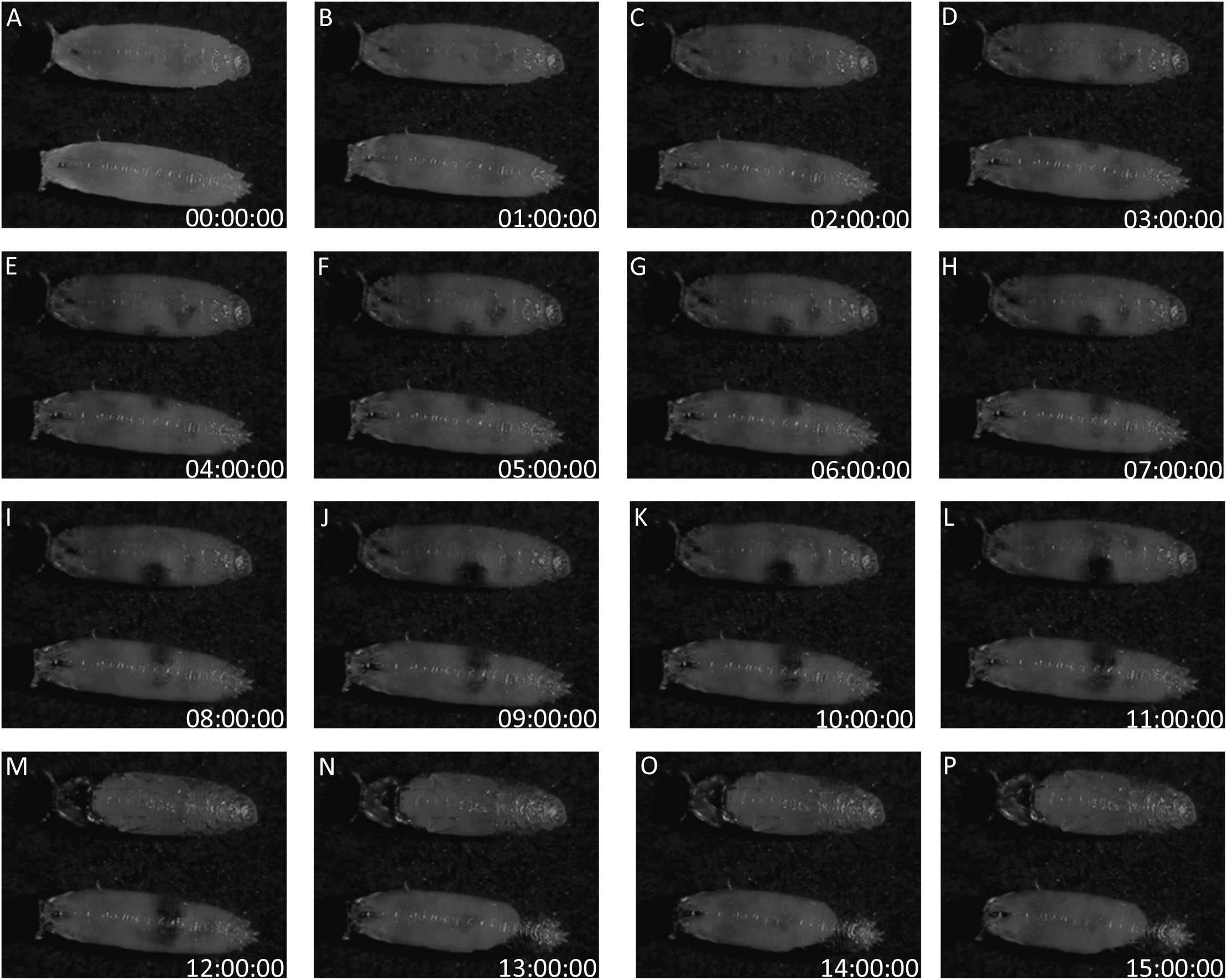
Air bubble movement fails in Nxt1 trans-heterozygote pupae. Still images of wild type (upper) and Nxt1 trans-heterozygotes (lower) for 15 hours. A) Faint visibility of air bubble in both genotypes. B-D) Air bubble becomes more prominent. E-G) Air bubble is clearly visible halfway along the pupa. I-L) Air bubble expands further. M) Air bubble has disappeared and permitted head eversion for wild type, whereas Nxt1 mutant still show air bubble. N) Larva body in Nxt1 mutant released from cuticle at posterior end and retracts to anterior. O-P) No further changes observed in both genotypes.

**Supplementary figure 2.**
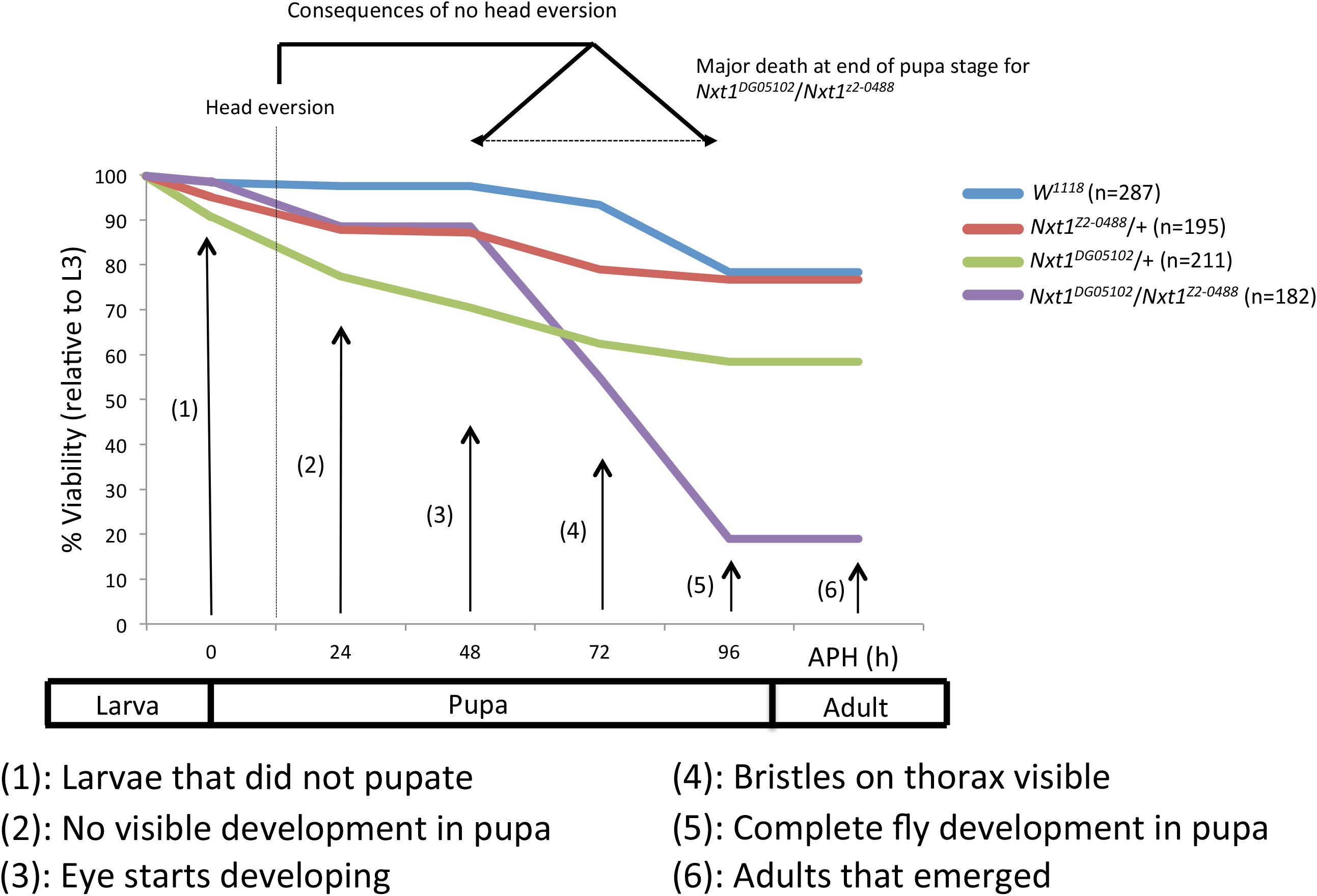
Timing of pupal lethality in Nxt1 trans-heterozygotes. Pupae were examined at six different time points. Many Nxt1 mutants had defects in head eversion, which becomes visible 48 hours into the pupa phase. Nxt1 hypomorph/+ heterozygotes showed a slight reduction in pupa viability.

**Supplementary figure 3.**
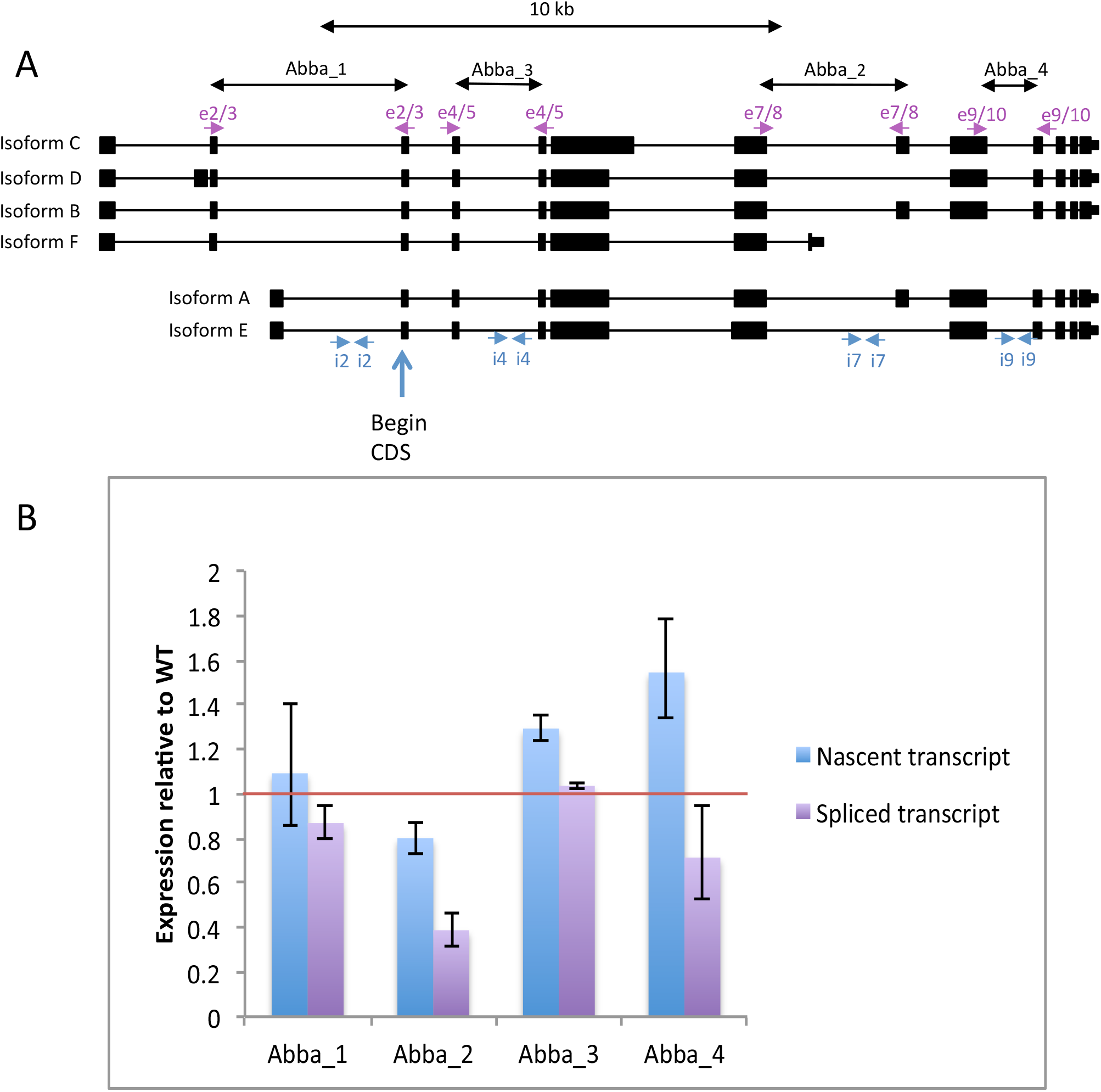
Reduced expression of spliced but not nascent *abba* transcripts in *Nxt1* trans-heterozygotes. A) Overview of six different *abba* isoforms, including the four regions used for nascent vs. spliced transcript analysis. B) qRT-PCR of four different regions in *abba* comparing nascent vs. spliced expression. *Rp49* was used for normalization.

**Supplementary table 1.** Known ecdysone responsive genes are not under-expressed in *Nxt1* trans-heterozygotes. Ecdysone responsive genes [21] were examined by looking their differential expression status in whole larval RNA-seq data from cuffdiff (P-value <0.05), a minimum fold change of 2 and a FPKM threshold of at least 10 FPKM in one of the two genotypes.

| Gene (wandering) | Differentially expressed? | Fold change (log2) | FPKM threshold? | Gene (stationary) | Differentially expressed? | Fold change (log2) | FPKM threshold? | Gene (prepupae) | Differentially expressed? | Fold change (log2) | FPKM threshold? |
| --- | --- | --- | --- | --- | --- | --- | --- | --- | --- | --- | --- |
| *Br* | NO | -0.10 | YES | *Br* | NO | -0.20 | YES | *Br* | NO | +0.02 | YES |
| *DHR3* | NO | -0.31 | NO | *DHR3* | NO | -0.87 | YES | *DHR3* | NO | -0.35 | YES |
| *EcR* | NO | -0.08 | YES | *EcR* | NO | +0.44 | YES | *EcR* | NO | +0.05 | YES |
| *Usp* | NO | +0.54 | YES | *Usp* | NO | +0.33 | YES | *Usp* | NO | -0.14 | YES |
| *Ecd* | NO | +0.28 | YES | *Ecd* | NO | +0.24 | YES | *Ecd* | NO | +0.36 | YES |
| *Edg78E* | NO | +0.64 | NO | *Edg78E* | NO | +2.15 | NO | *Edg78E* | NO | +1.05 | NO |
| *Edg84A* | NO | - | - | *Edg84A* | NO | - | - | *Edg84A* | NO | - | - |
| *Edg91* | NO | -0.05 | YES | *Edg91* | NO | +0.46 | YES | *Edg91* | NO | -0.16 | NO |
| *Eip55E* | NO | +0.38 | YES | *Eip55E* | NO | +0.30 | YES | *Eip55E* | NO | 0.00 | YES |
| *Eip63E* | NO | 0.00 | YES | *Eip63E* | NO | -0.04 | YES | *Eip63E* | NO | +0.36 | YES |
| *Eip63F-1* | NO | -0.74 | YES | *Eip63F-1* | YES | -0.97 | YES | *Eip63F-1* | YES | +1.00 | YES |
| *Eip63F-2* | NO | +0.26 | NO | *Eip63F-2* | NO | +0.06 | NO | *Eip63F-2* | NO | +0.85 | NO |
| *Eip71CD* | YES | -0.87 | YES | *Eip71CD* | NO | -0.47 | YES | *Eip71CD* | NO | -0.18 | YES |
| *Eip74EF* | NO | +0.03 | YES | *Eip74EF* | NO | 0.00 | YES | *Eip74EF* | NO | +0.42 | YES |
| *Eip75B* | NO | -0.37 | YES | *Eip75B* | NO | -0.49 | YES | *Eip75B* | NO | -0.30 | YES |
| *Eip78C* | NO | -0.31 | YES | *Eip78C* | NO | -0.37 | YES | *Eip78C* | YES | +0.68 | YES |
| *Eip93F* | NO | +0.35 | NO | *Eip93F* | NO | -0.43 | YES | *Eip93F* | NO | +0.29 | YES |
| *Ftz-f1* | NO | +0.46 | NO | *Ftz-f1* | NO | +0.66 | NO | *Ftz-f1* | NO | -0.47 | NO |
| *Eig71Ea* | NO | -1.18 | YES | *Eig71Ea* | YES | -0.57 | YES | *Eig71Ea* | NO | +0.32 | YES |
| *Eig71Eb* | NO | -1.16 | YES | *Eig71Eb* | YES | -1.62 | YES | *Eig71Eb* | YES | -0.78 | YES |
| *Eig71Ec* | NO | -2.16 | NO | *Eig71Ec* | NO | -0.73 | YES | *Eig71Ec* | NO | 0.00 | YES |
| *Eig71Ed* | NO | -1.27 | YES | *Eig71Ed* | YES | -0.93 | YES | *Eig71Ed* | YES | -0.42 | YES |
| *Eig71Ee* | NO | -0.28 | YES | *Eig71Ee* | NO | -0.08 | YES | *Eig71Ee* | YES | +0.85 | YES |
| *Eig71Ef* | NO | -2.44 | NO | *Eig71Ef* | YES | -1.14 | YES | *Eig71Ef* | NO | -0.34 | YES |
| *Eig71Eg* | NO | -1.99 | YES | *Eig71Eg* | YES | -1.55 | YES | *Eig71Eg* | YES | -0.97 | YES |
| *Eig71Eh* | NO | - | - | *Eig71Eh* | NO | -3.34 | NO | *Eig71Eh* | YES | -1.48 | YES |
| *Eig71Ei* | NO | - | - | *Eig71Ei* | NO | -2.50 | NO | *Eig71Ei* | NO | -0.75 | YES |
| *Eig71Ej* | NO | - | - | *Eig71Ej* | NO | - | - | *Eig71Ej* | NO | -0.92 | NO |
| *Eig71Ek* | NO | - | - | *Eig71Ek* | NO | - | - | *Eig71Ek* | NO | +1.21 | NO |

**Supplementary table 2.**
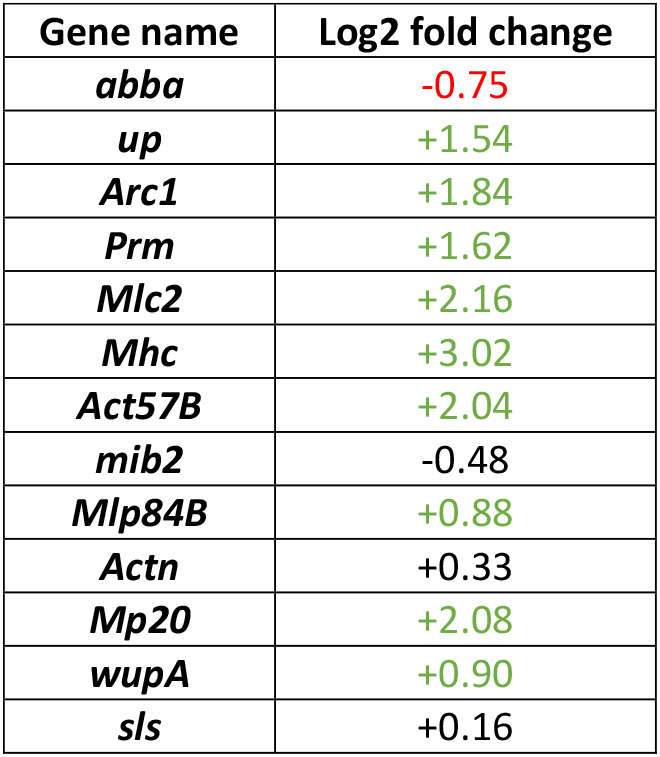
Overview of larval muscle genes from whole larval RNA sequencing. Thirteen larval muscle genes with log2 fold change from RNA sequencing showing majority of genes up regulated with only one down regulated. Red = more than 1.5-fold down regulated. Green = more than 1.5-fold up regulated.

